# Deciphering the altered conformational states of bifunctional thaumarchaeal crotonyl-CoA hydratase and 3-hydroxypropionyl-CoA dehydratase from *Nitrosopumilus maritimus*

**DOI:** 10.1101/2025.07.07.663511

**Authors:** Ebru Destan, Jungmin Kang, Takehiko Tosha, Makina Yabashi, İlkin Yapıcı, Bradley B. Tolar, Cahine Kulakman, Zelis Nergiz, Hiroaki Matsuura, Yoshiaki Kawano, Samuel Deutsch, Yasuo Yoshikuni, Christopher A. Francis, Soichi Wakatsuki, Hasan DeMirci

## Abstract

The thaumarchaeal 3-hydroxypropionate/4-hydroxybutyrate (3HP/4HB) cycle represents one of the most efficient mechanisms for CO2 fixation discovered to date. Within this cycle, the enzyme encoded by Nmar_1308 from *Nitrosopumilus maritimus* SCM1 plays a crucial role due to its dual functionality as both a crotonyl-CoA hydratase (CCAH) and a 3-hydroxypropionyl-CoA dehydratase (3HPD). Although the importance of a bifunctional enzyme for lowering the cost of biosynthesis, the details of structural dynamics are still missing. Here, in addition to our cryogenic temperature structures, we determined the first ambient temperature structures of the Nmar_1308 protein by Serial Femtosecond X-ray Crystallography (SFX). The determined structures capture previously unobserved conformational dynamics of the Nmar_1308 protein, providing invaluable information for future synthetic biology applications.

## Introduction

Carbon exchange is crucial in preserving the earth’s ecological balance and CO_2_ fixation, which transforms inorganic CO_2_ into organic compounds, plays a vital role in this process. CO_2_ fixation is particularly significant for autotrophic organisms, which utilize the 3-hydroxypropionate/4-hydroxybutyrate (3HP/4HB) cycle.^**[1]**^ This cycle accounts for approximately 1% of global CO_2_ fixation.^**[2]**^ Given the increasing levels of atmospheric CO_2_ due to human activities, understanding the natural CO_2_ fixation mechanisms is more relevant than ever, as they offer insights into potential biotechnological applications for carbon sequestration. The ability of autotrophic organisms to thrive in environments with low energy availability underscores the importance of their carbon and nitrogen fixation strategies, which have evolved to maximize efficiency in resource-limited conditions.^**[3]**^ This efficiency is not only crucial for their survival but also for their substantial contribution to the Earth’s biogeochemical cycles. ^**[4]**^

A unique phylum of Archaea, namely Thaumarchaeota, plays a significant role in global carbon and nitrogen cycling.^**[5]**^ As Thaumarchaeota can thrive in nutrient-poor habitats, they are widely distributed across diverse environments from the deep sea to terrestrial soils.^**[6]**^ One of the defining characteristics of Thaumarchaeota, including the species *Nitrosopumilus maritimus* (*N. maritimus)*, is their ability to oxidize ammonia to nitrite, an essential step in the nitrogen cycle.^**[7,8]**^ This capacity is particularly important in marine environments, where *N. maritimus* has been shown to dominate ammonia oxidation, contributing substantially to global nitrogen turnover.^**[9]**^ The ecological success of Thaumarchaeota, particularly in low-nutrient environments, is largely attributed to their highly efficient metabolic pathways, such as the 3HP/4HB cycle for carbon fixation.^**[1,2]**^ In this cycle, one acetyl-CoA and two bicarbonate molecules are first converted to succinyl-CoA through a 3-hydroxypropionate intermediate.^**[1]**^ The second half of the cycle involves the conversion of succinyl-CoA, regenerating acetyl-CoA, and producing two molecules of acetyl-CoA via a 4-hydroxybutyrate intermediate.^**[10]**^ One of these acetyl-CoA molecules is then used as a carbon source for the subsequent cycle.^**[11]**^ The 3HP/4HB cycle is one of the most energy-efficient carbon fixation pathways known, requiring fewer ATP molecules compared to other autotrophic pathways like the Calvin-Benson-Bassham cycle.^**[1]**^ The 3HP/4HB cycle is unique to certain Archaea and allows *N. maritimus* to fix CO_2_ under aerobic conditions, converting inorganic carbon into organic molecules necessary for growth and survival **(Figure 1A)**.^**[1,12]**^

**Figure 1:**
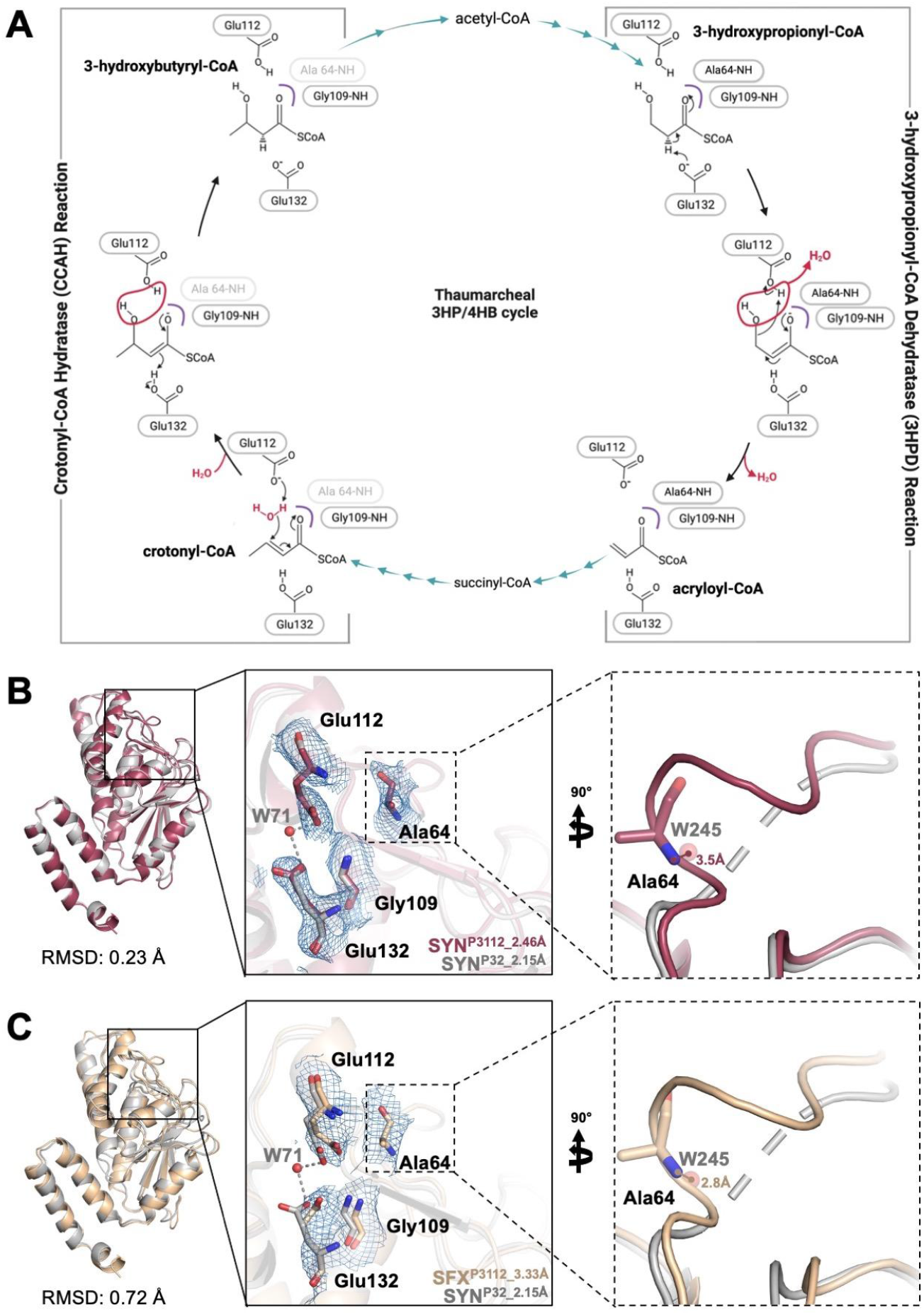
Representation of biofunctional enzymes from *Nitrosopumilus maritimus*. **(A)** Hydratase/Dehydratase reactions are illustrated in the thaumarcheal 3HP/4HB cycle. **(B)** SYN^P3112_2.46Å^ structure is superposed with the previously published Nmar_1308 protein structure (SYN^P32_2.15Å^; PDB ID: 7EUM). **(C)** SFX^P312_3.33Å^ structure is superposed with SYN^P32_2.15Å^ structure.

Previously, the structure of two bifunctional enzymes in this cycle was obtained from different organisms. The first one was obtained from *Metallosphaera sedula* (5ZA1; Msed_2001), and the most recent one from *N. maritimus* SCM1 (7EUM;

Nmar_1308).^**[13,14]**^ Both these enzymes play a bifunctional role during crotonyl-CoA hydratase (CCAH) and 3-hydroxypropionyl-CoA dehydratase (3HPD) reactions. As a focus of this study, Nmar_1308 protein acts both as a 3HPD and a CCAH.^**[1,15]**^ In its capacity as a 3HPD, Nmar_1308 catalyzes the conversion of 3-hydroxypropionyl-CoA to acryloyl-CoA and water. Conversely, as a CCAH, Nmar_1308 converts crotonyl-CoA to 3-hydroxybutyryl-CoA through a dehydration reaction. The versatility of Nmar_1308, in addition to the role of another bifunctional enzyme, acetyl-CoA/propionyl-CoA carboxylase, in the *N. maritimus* 3HP/4HB cycle enhances metabolic efficiency by reducing the overall demand for protein synthesis.^**[1]**^ This functional flexibility contributes to streamlining biosynthetic costs, improving the efficiency of energy and resource usage, and optimizing cellular processes.

Here, together with our cryogenic temperature structures, we determined the first ambient temperature structure of the Nmar_1308 protein. These structures revealed the previously unobserved conformational dynamics of the bifunctional enzyme, highlighting the missing details of structural dynamics during hydratase/dehydratase reactions in the thaumarchaeal 3HP/4HB cycle.

## Results and discussions

### Cryogenic and ambient temperature structure of apo-form Nmar_1308 protein

The six apo-form structures of Nmar_1308 protein were determined in trimeric and hexameric forms. The trimeric structures include SYN^P3112_2.46Å^ in the P3112 space group with 2.46 Å resolution, SYN^P32_2.71Å^ in the P32 space group with 2.71Å resolution, SYN^P3212_3.29Å^ in the P3212 space group with 3.29Å resolution, SFX^P3112_3.33Å^ in the P3112 space group, and SFX ^P3212_3.40Å^ in the P3212 space group with 3.40 Å resolution **(Table 1)**. The hexameric form is represented by SYN^C121_2.96Å^, determined in the C121 space group with 2.96Å resolution **(Table 1)**. The P3112, P32, and P3212 space groups’ crystals from the hexagonal-trigonal lattice appeared more compact and homogeneous, which helped to increase symmetry. The elongated and less symmetrical monoclinic lattice crystals (C121 space group), on the other hand, suggested that the protein’s conformation might be less stable.^**[16]**^ The difference in the space groups was reflected in crystal morphology as well **(Figure S1-2, Supporting Information)**.

**Table 1:** Data collection and refinement statistics. ^1^ CC^*^ value is presented for the dataset. ^2^ CC (*1/2*) value is presented for the dataset.

| Datasets | SYN<sup>P3112_2.46Å</sup> | SYN<sup>P32_2.71Å</sup> | SYN<sup>C121_2.96Å</sup> | SYN<sup>P3212_3.29Å</sup> | SFX<sup>P3112_3.33Å</sup> | SFX<sup>P3212_3.40Å</sup> |
| --- | --- | --- | --- | --- | --- | --- |
| PDB IDs | 9UL8 | 9UL9 | 9ULA | 9ULB | 9ULC | 9ULD |
| Data collection temperature | Cryogenic | Cryogenic | Cryogenic | Cryogenic | Ambient | Ambient |
| Data collection |  |  |  |  |  |  |
| X-ray source | Spring-8 (BL32XU) | Spring-8 (BL32XU) | Spring-8 (BL32XU) | Spring-8 (BL32XU) | SACLA (BL2) | SACLA (BL2) |
| Space group | P3112 | P32 | C121 | P3212 | P3112 | P3212 |
| a, b, c (Å) | 75.97, 75.97, 209.42 | 74.19, 74.19, 206.85 | 356.96, 130.53, 82.64 | 76.3, 76.3, 210.11 | 81.13, 81.13, 216.35 | 81.33, 81.33, 219.87 |
| α, β, γ (°) | 90.00, 90.00, 120.00 | 90.00, 90.00, 120.00 | 90, 100.174, 90 | 90.00, 90.00, 120.00 | 90.00, 90.00, 120.00 | 90.00, 90.00, 120.00 |
| Resolution (Å) | 37.99-2.46 (2.52-2.46) | 40.29-2.71 (2.78-2.71) | 43.92-2.96 (3.03-2.96) | 41.12-3.29 (3.44-3.29) | 32.46-3.33 (3.47-3.33) | 32.71-3.40 (3.82-3.40) |
| I / σI | 10.17 (0.66) | 8.91 (1.51) | 8.85 (1.12) | 6.06 (0.95) | 4.23 (1.95) | 3.70 (1.73) |
| <sup>1</sup>CC* or <sup>2</sup>CC(1/2) | <sup>2</sup>99.2 (50.1) | <sup>2</sup>96.8 (62.2) | <sup>2</sup>98.9 (58.3) | <sup>2</sup>98.4 (45.00) | <sup>1</sup>0.97 (0.86) | <sup>1</sup>0.96 (0.76) |
| Completeness (%) | 100.00 (100.00) | 100.00 (100.00) | 99.99 (100.00) | 100.00 (99.99) | 100.00 (100.00) | 100.00 (100.00) |
| Redundancy | 96.99 (83.67) | 16.37 (11.76) | 11.83 (10.51) | 13.19 (11.76) | 69.84 (64.5) | 62.81 (59.5) |
| No. unique reflections | 25601 (4123) | 34665 (5506) | 77784 (12533) | 10985 (1754) | 55194 (2748) | 56420 (6425) |
| No. observed reflections | 2482940 (344979) | 567622 (64760) | 920614 (131766) | 144927 (20637) | 3854822 (177246) | 3543998 (382287) |
| Refinement |  |  |  |  |  |  |
| Resolution (Å) | 37.99-2.46 (2.52-2.46) | 40.29-2.71 (2.78-2.71) | 43.91-2.96 (3.03-2.96) | 41.12-3.29 (3.44-3.29) | 24.04-3.33 (3.54-3.33) | 24.43-3.40 (3.61-3.40) |
| R<sub>work</sub> / R<sub>free</sub> | 0.37/0.40 (0.40/0.40) | 0.37/0.39 (0.35/0.40) | 0.44/0.46 (0.40/0.41) | 0.35/0.41 (0.33/0.40) | 0.32/0.37 (0.31/0.32) | 0.32/0.36 (0.34/0.37) |
| No. atoms |  |  |  |  |  |  |
| Protein | 5491 | 5481 | 10832 | 5260 | 5473 | 5473 |
| Ligand/ion/ Water | 35 | 62 | 48 | 7 | - | - |
| B-factors (Å<sup>2</sup>) |  |  |  |  |  |  |
| Protein | 45.42 | 36.82 | 50.88 | 72.60 | 54.49 | 63.66 |
| Ligand/ion/ Water | 37.81 | 24.36 | 46.88 | 32.04 | - | - |
| R.m.s. deviations |  |  |  |  |  |  |
| Bond lengths (Å) | 0.005 | 0.002 | 0.003 | 0.004 | 0.003 | 0.002 |
| Bond angles (°) | 0.904 | 0.476 | 0.619 | 0.750 | 0.679 | 0.588 |
| Ramachandran plot |  |  |  |  |  |  |
| Favored (%) | 92.56 | 94.21 | 94.42 | 93.10 | 90.77 | 90.50 |
| Allowed (%) | 7.44 | 5.79 | 5.58 | 6.31 | 8.82 | 9.37 |
| Disallowed (%) | 0.00 | 0.00 | 0.00 | 0.59 | 0.41 | 0.14 |

To highlight the key conformational changes, the SFX^P3112_3.33Å^ and SYN^P3112_2.46Å^ structures were superposed with the previously available cryogenic structure of Nmar_1308 protein (SYN^P32_2.15Å^; PDB ID: 7EUM) **(Figure 1B,C)**.^**[14]**^ The SYN^P3112_2.46Å^ structure and SYN^P32_2.15Å^ have a very low root mean square deviation (RMSD) (0.23 Å), which suggests a high degree of structural similarity **(Figure 1B; Table 2)**. However, the comparison of SFX^P3112_3.33Å^ and SYN^P32_2.15Å^ structures indicated more substantial atomic position deviations with a slightly higher RMSD (0.72 Å) **(Figure 1C; Table 2)**. To further investigate differences in protein stability, b-factor analysis was performed by using Serial Femtosecond Crystallography (SFX) and synchrotron radiation (SYN) structures. B-factor analysis revealed that the active site of the new structures appears to be more rigid and well-defined compared to SYN^P32_2.15Å^ **(Figure 2; Figure S3, Supporting Information)**. Particularly in the active site region, catalytic residues (Glu112, Gly109, Ala64, and Glu132) remained well-ordered. The cryogenic temperature structures SYN^P3112_2.46Å^, SYN^P32_2.71Å^, and SYN^C121_2.96Å^ exhibited low b-factors across the protein, indicating a stable conformation. In contrast, the SYN^P3212_3.29Å^ structure demonstrated higher B-factors, suggesting increased flexibility **(Figure S3B, Supporting Information)**. For the ambient temperature structures, the SFX ^P3212_3.40Å^ structure was observed as slightly more flexible compared to the SFX^P3112_3.33Å^ structure. It suggests that the SFX^P3112_3.33Å^ structure might represent a different, more stable conformational state.

**Table 2:** RMSD(Å) values for Chain A in **Figure 3**.

| **RMSD (Å)**<br>**_Chain A** | SYN<sup>P3112_2.46Å</sup> | SYN<sup>P32_2.71Å</sup> | SYN<sup>C121_2.96Å</sup> | SYN<sup>P3212_3.29Å</sup> | SFX<sup>P3112_3.33Å</sup> | SFX<sup>P3212_3.40Å</sup> | 7EUM |
| --- | --- | --- | --- | --- | --- | --- | --- |
| SYN<sup>P3112_2.46Å</sup> | 0 | 0.25 | 0.33 | 0.40 | 0.71 | 0.80 | 0.23 |
| SYN<sup>P32_2.71Å</sup> | 0.25 | 0 | 0.24 | 0.37 | 0.74 | 0.84 | 0.23 |
| SYN<sup>C121_2.96Å</sup> | 0.33 | 0.24 | 0 | 0.45 | 0.71 | 0.78 | 0.32 |
| SYN<sup>P3212_3.29Å</sup> | 0.40 | 0.37 | 0.45 | 0 | 0.76 | 0.88 | 0.35 |
| SFX<sup>P3112_3.33Å</sup> | 0.71 | 0.74 | 0.71 | 0.76 | 0 | 0.81 | 0.72 |
| SFX<sup>P3212_3.40Å</sup> | 0.80 | 0.84 | 0.78 | 0.88 | 0.81 | 0 | 0.83 |
| 7EUM | 0.23 | 0.23 | 0.32 | 0.35 | 0.72 | 0.83 | 0 |

**Figure 2:**
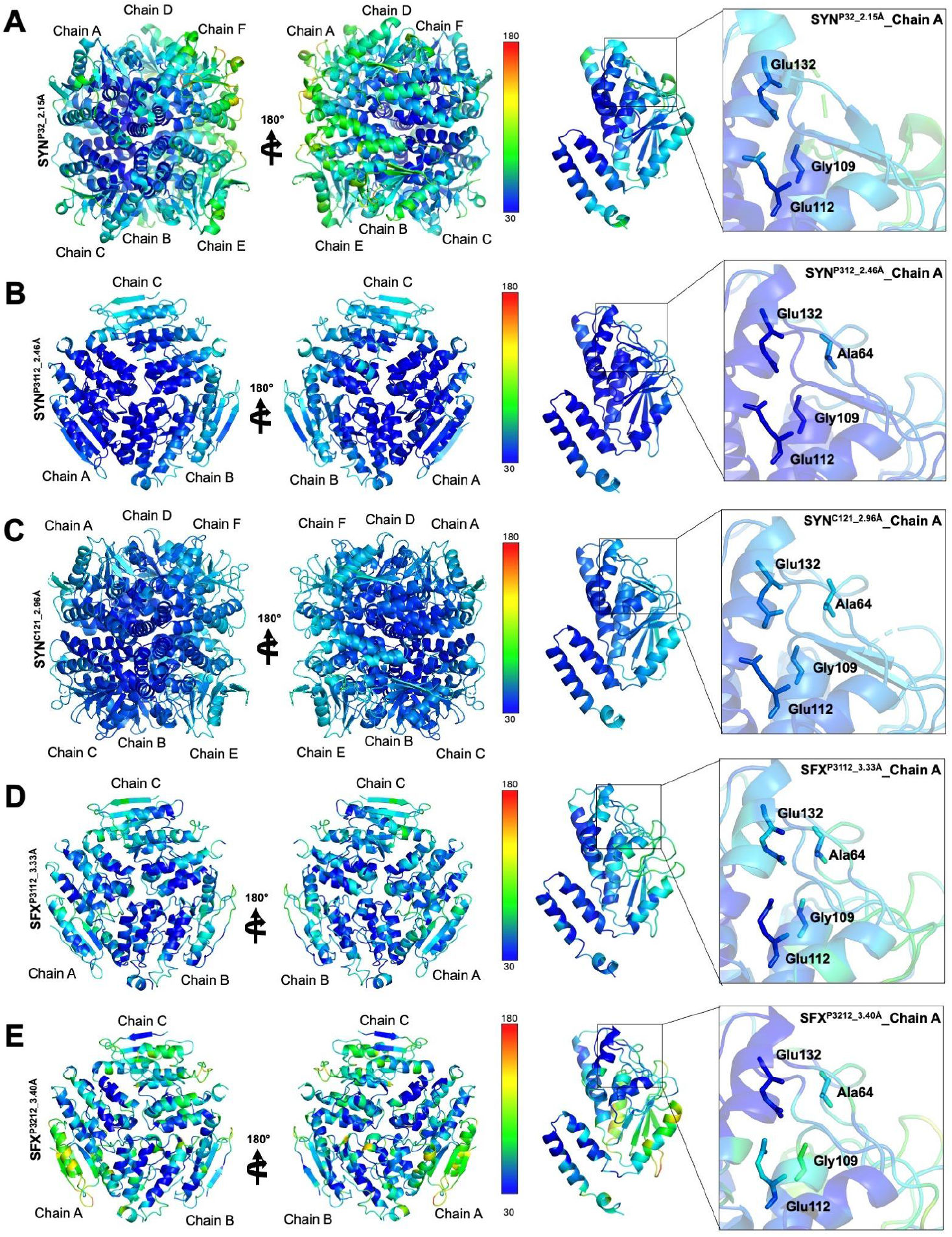
Apo-form Nmar_1308 protein crystal structures based on B factor. **(A-E)** All structures are colored based on the B-factor (Spectrum range: 30 to 180 Å^2^). Chain A of each structure is represented to compare the stability around the substrate binding pocket.

### Conformational changes at the active site

The structural comparisons presented in Figure 3, along with the RMSD values from Table 2, offer key insights into the dynamic nature of the substrate-binding pocket. The superposition of the SFX^P3112_3.33Å^ and SFX^P3212_3.40Å^ structures revealed minimal side-chain displacements in residues Glu132, Ala64, Gly109, and Glu112 (**Figure 3A**). Although structural conservation at the active site, two ambient temperatures were superposed with an RMSD value of 0.81 Å, which is slightly higher compared to other pairs in Table 2. These originate from the flexible loop regions, whose conformation slightly differs between the two structures. The comparison with the previously available cryogenic temperature structure of Nmar_1308 (SYN^P32_2.15Å^; PDB ID: 7EUM) revealed a significant conformational change at the active site. Additionally, the superposition of SFX^P3112_3.33Å^ and SYN^P3112_2.46Å^ structures indicated significant conformational changes, particularly in the position of Glu112 and Glu132 (**Figure 3B**). This indicates moderate flexibility in response to changes in experimental conditions, as supported by an increase in the RMSD values. These adjustments suggest that the protein undergoes conformational changes under varying temperatures. A comparison of SFX^P3112_3.33Å^ and SYN^P32_2.71Å^ indicates major conformational changes in the position of Glu 112 and Glu132, while minor conformational changes for Ala 64 and Gly109 (**Figure 3C**). Here, it was observed that cryogenic temperature structures are more structurally similar to the SYN^P32_2.15Å^ structure. In these circumstances, our SFX structures provide previously unobserved structural dynamics at ambient temperature. Additionally, the superposition of SFX^P3112_3.33Å^ and SYN^C121_2.96Å^ reveals significant displacements for catalytic residues, indicating a different conformational state in the C121 space group (**Figure 3D**). Similar to other cryogenic temperature structures, the alignment of SFX^P3112_3.33Å^ and SYN^P3212_3.29Å^ demonstrated conformational changes on key residues such as Glu112, and minor conformational changes on Ala 64, Gly 109, and Glu 132 (**Figure 3E**). Together with the alignment of all cryogenic structures **(Figure 3F)**, it was supported that there are only minor conformational changes, except in the SYN^C121_2.96Å^ structure. This indicates a different conformational state, which may have originated due to different crystal morphology.

**Figure 3:**
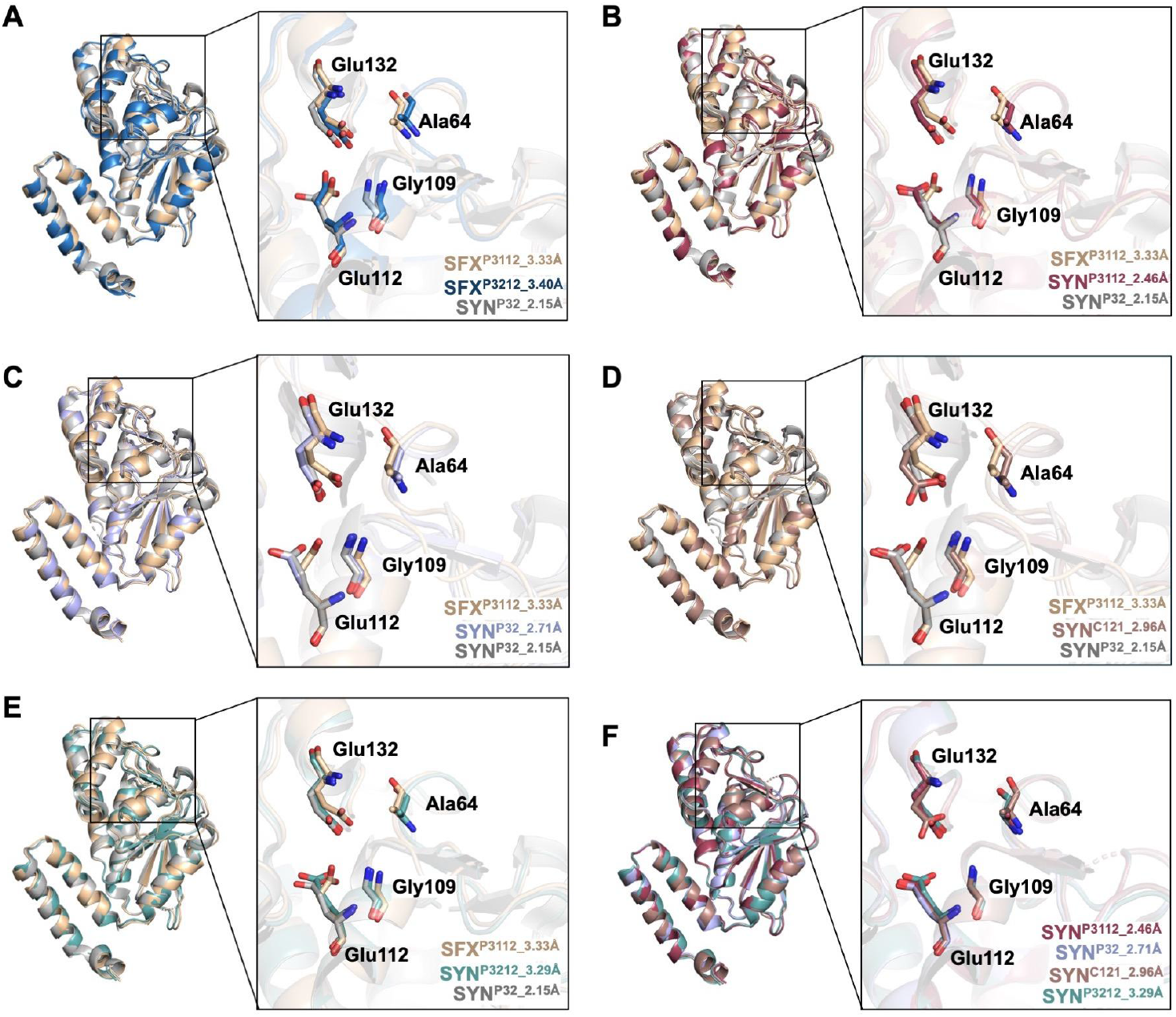
Comparison of cryogenic and ambient temperature apo-form Nmar_1308 structures. **(A)** Ambient temperature structures (SFX^P312_3.33Å^ and SFX^P312_3.40Å^) are superposed with the previously available cryogenic Nmar_1308 protein structure (SYN^P32_2.15Å^; PDB ID: 7EUM). **(B)** SFX^P312_3.33Å^ is superposed with SYN^P312_2.46Å^ and SYN^P32_2.15Å^ structures. **(C)** SFX^P312_3.33Å^ is superposed with SYN^P31_2.71Å^ and SYN^P32_2.15Å^ structures. **(D)** SFX^P312_3.33Å^ is superposed with SYN^C121_2.96Å^ and SYN^P32_2.15Å^ structures. **(E)** SFX^P312_3.33Å^ is superposed with SYN^P312_3.29Å^ and SYN^P32_2.15Å^ structures. **(F)** All synchrotron structures are superposed to observe conformational changes in the active site at cryogenic temperature. RMSD(Å) values for Chain A are indicated in Table 2.

These observed conformational changes may highlight functional flexibility in the protein’s active or binding sites. The RMSD values presented in **Table 2** quantitatively support these structural observations. Overall, the results from Figure 3 and the RMSD analysis in Table 2 reveal a balance between structural stability and flexibility within the protein. While some regions, such as those around Ala64, remain highly conserved across different conditions, other areas exhibit conformational adjustments, particularly under distinct crystallization environments. These shifts likely contribute to the protein’s ability to adapt its structure for functional purposes, such as substrate binding or catalytic activity.

### New insight into the previously disordered region

The revealed Nmar_1308 protein structures are in apo-form, and a detailed examination of their ligand-binding pocket is essential to understanding the binding mode. The hydratase from *Rattus norvegicus* (PDB ID: 1DUB) with its ligand acetoacetyl-CoA, dehydratase from *Myxococcus xanthus* (PDB ID: 5JBX) with its ligand CoA, and bifunctional hydratase-dehydratase from *Metallosphaera sedula* (PDB ID: 5ZAI) were superposed with the apo-form Nmar_1308 protein structures to closely investigate important aspects of the active site **(Figure 4A,C,E; Figures S4-8, Supporting Information)**. Superposition analysis revealed structural similarities and differences, which provide insights into the altered binding pocket and substrate interactions. While the active site residues of cryogenic temperature structures show only slight changes compared to homologous holo-structures, the ambient temperature structures (SFX^P3112_3.33^ and SFX^P3212_3.40Å^) exhibit significantly altered conformations of key residues and shifts in their positions along the main chain **(Figure 4B,D,F; Figure S7, Supporting Information**). These differences suggest potential temperature-dependent flexibility, which may have implications for the structural dynamics under physiological conditions. Among the key residues involved in stabilizing the substrate binding pocket are Gly109, Glu112, Glu132, and the dynamic Ala64. Ala64, located at the active site, undergoes radical conformational shifts, especially in the presence of a substrate, as shown in the superposed holo-structures. These changes suggest significant structural rearrangements in the active site upon ligand binding, possibly enhancing catalytic efficiency or substrate affinity **(Figure 4; Figures S4-7, Supporting Information)**.

**Figure 4:**
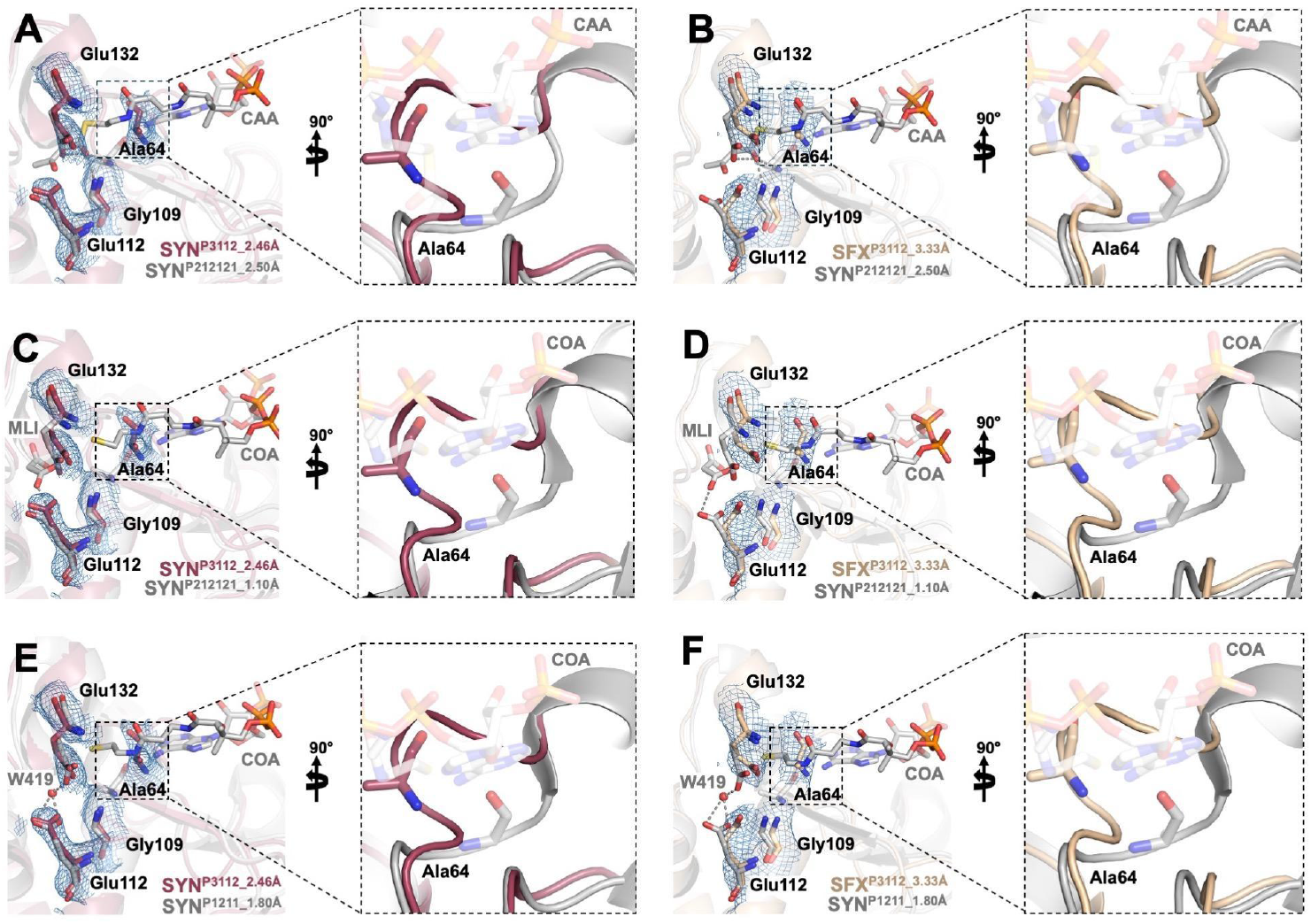
Comparison of apo-form Nmar_1308 protein structures with previously available homologous hydratase/dehydratase structures. **(A,C,E)** To compare the substrate binding pocket, the SYN^P3112_2.46Å^ structure is superposed with a *Rattus norvegicus* hydratase structure (SYN^P212121_2.50Å^; PDB ID: 1DUB), a *Myxococcus xanthus* dehydratase structure (SYN^P212121_1.10Å^; PDB ID: 5JBX), and another biofunctional enzyme from *Metallosphaera sedula* (SYN^P1211_1.80Å^; PDB ID: 5ZAI). **(B,D,F)** SFX^P312_3.33Å^ structure is superposed with a hydratase structure (SYN^P212121_2.50Å^; PDB ID: 1DUB), dehydratase structure (SYN^P212121_1.10Å^; PDB ID: 5JBX), and another biofunctional enzyme from *Metallosphaera sedula* (SYN^P1211_1.80Å^; PDB ID: 5ZAI).

As previously proposed, Ala64 and Gly109 form an oxyanion hole that stabilizes the negative charge on the substrate’s carbonyl oxygen, a key step in the catalytic process.^**[14]**^ In our prior studies, Ala64 was shown to lack a defined electron density, indicating flexibility in this region (SYN^P32_2.15Å^; PDB ID: 7EUM)^**[14]**^. However, in the newly determined apo-form Nmar_1308 structures, the disordered residues 62-68, forming a surface-exposed loop, now exhibit well-defined electron density **(Figure 5)**. This clearer representation suggests that the flexibility of this loop may be important in the protein’s function, potentially modulating access to the active site during substrate binding or product release. Moreover, the highest resolution structure, SYNP^3112_2.46Å^, reveals the positions of side chains with alternative conformations compared to other structures, shedding light on potential flexibility and structural rearrangements within the active site **(Figure 5A)**. This newly observed flexibility of the side chains further supports the hypothesis that these dynamic rearrangements play a critical role in substrate stabilization and binding affinity, potentially allowing for fine-tuned interactions with various ligands. Further to highlight the mechanism behind the oxyanion hole formation, the active site of SYN^P3112_2.46Å^ and SFX^P312_3.33Å^ structures were superposed with a hydratase SYN^P212121_2.50Å^ structure **(Figure 6)**. The distances between the carbonyl oxygen (O1) of a substrate and the backbone NH groups of amino acids (Ala64 and Gly109) are crucial for the proper formation of the oxyanion hole and stabilization of the substrate. It is observed that the distances with Ala64-NH are substantially longer for both our apo cryogenic and ambient temperature structures (4.9Å and 4.4Å, respectively) than those observed in the substrate bound hydratase, 3.3Å (SYN^P212121_2.50Å^), revealing the altered dynamics of the active site with the entrance of the substrate **(Figure 6A,B)**. Together with the displacement of the loop, the distance for the side-chain of critical residues (Glu112 and Glu132) is decreasing for the apo structures **(Figure 6C-F)**. These findings indicate that the active site undergoes conformational changes to form a proper oxyanion hole during/after the substrate binding and stabilize the substrate during the hydratase/dehydratase reaction.

**Figure 5:**
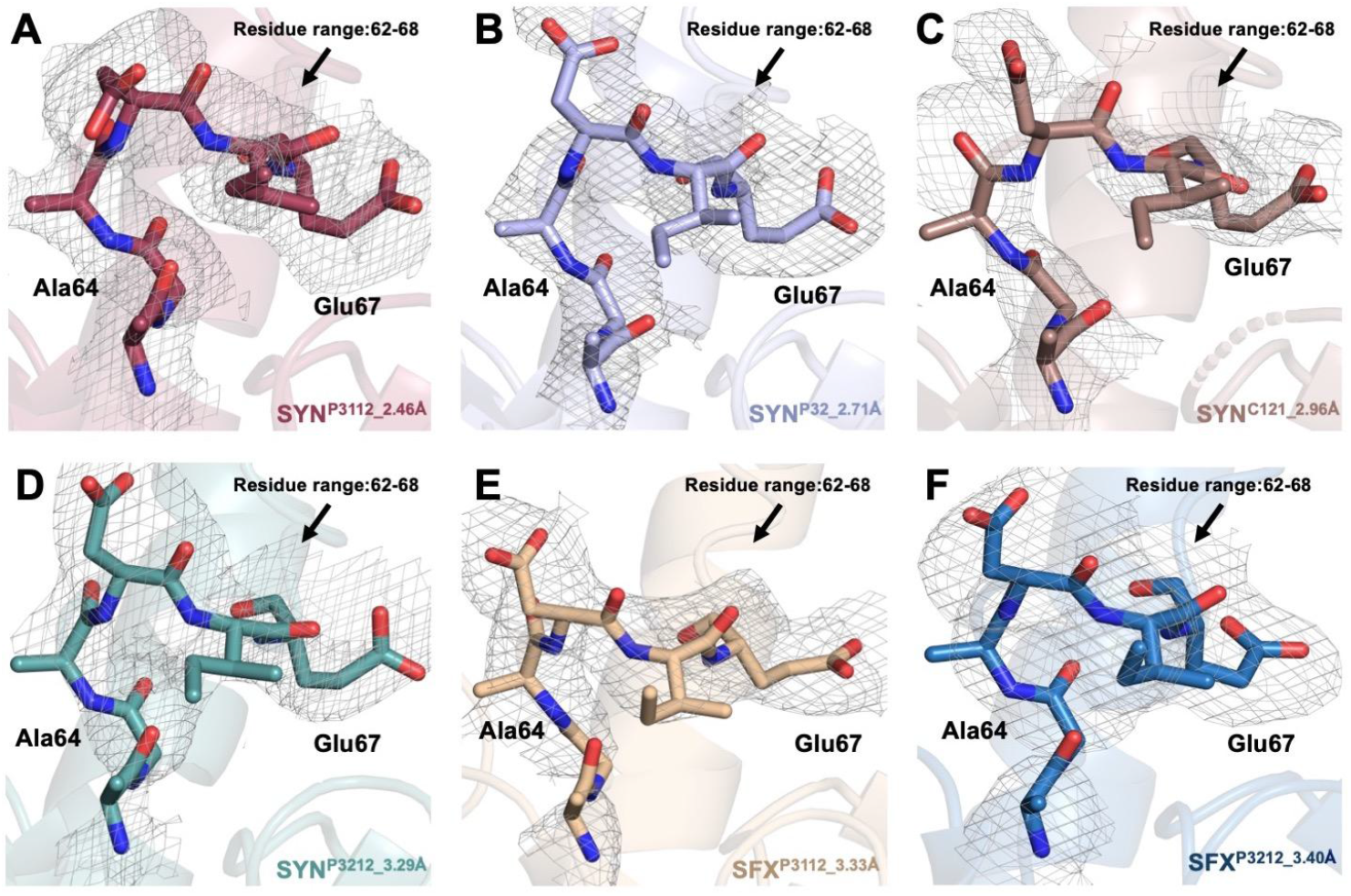
2Fo-Fc electron density map of the loop region in the substrate binding pocket. **(A-F)** The residues between Ala62 and Glu67, which were previously detected as disordered in the Nmar_1308 protein (PDB ID 7EUM), are indicated by stick representation. 2Fo-Fc electron density map is contoured at the 1σ level and colored in gray.

**Figure 6:**
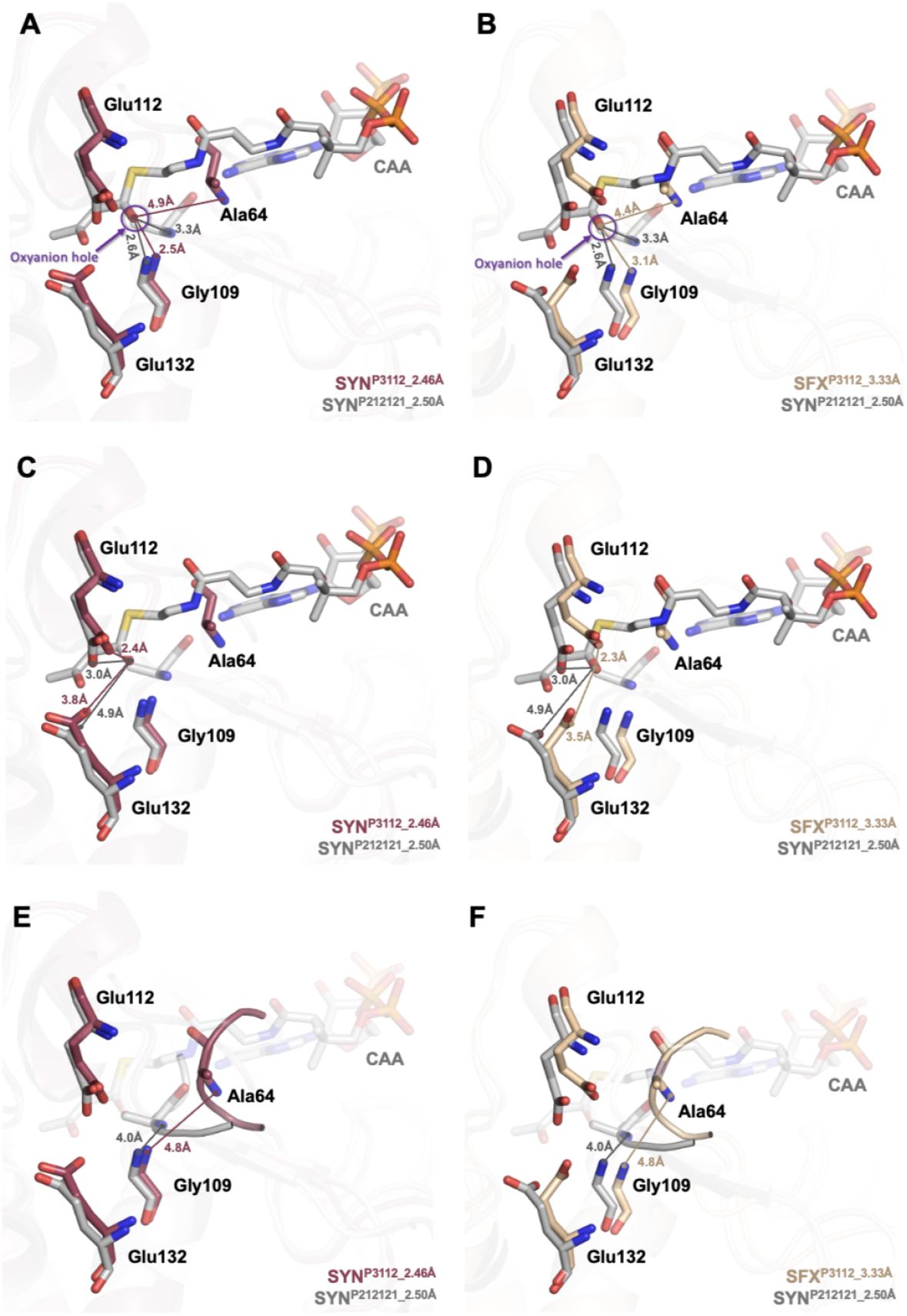
Representation of oxyanion hole formation during the hydratase/dehydratase reaction. SYN^P3112_2.46Å^ and SFX^P312_3.33Å^ structures were superposed with a hydratase structure (SYNP212121_2.50Å; PDB ID: 1DUB) to highlight the details of enzyme activity. **(A,B)** The distances between the carbonyl oxygen (O1) of a substrate and Ala64-NH/Gly109-NH are calculated for both ambient and cryogenic temperature structures. **(C,D)** The distances between the carbonyl oxygen (O1) of a substrate and Glu112-OE2/Glu132-OE1 are calculated for both ambient and cryogenic temperature structures to indicate the side chain displacements for critical residues. **(E-F)** The distances between Ala64-NH and Gly109-NH are calculated for both ambient and cryogenic temperature structures to indicate the loop displacement in the presence of the acetoacetyl-coenzyme A (CAA).

### Hydrogen bond network and the function of Trp141 at the active site

Detailed analysis of interactions on catalytic residues (Glu112, Glu132) at the active site may provide a better understanding of how a bifunctional enzyme functions **(Figure 7; Figure S9, Supporting Information)**. Compared to the previously published structure of apo-form Nmar_1308 protein (SYN^P32_2.15Å^; PDB ID: 7EUM), here, we observed a different conformational state where the critical water (W71) molecule was removed during 3HPDD reaction **(Figure 1A)**.^**[14]**^ In the SYN^P3112_2.46Å^ structure, Glu112 interacts with the residues Gly108, Ser116, Gly143, and Trp141 while Glu132 interacts with the residue Glu130 **(Figure 7D)**. The same interactions were observed for SYN^P32_2.71Å^, SYN^C121_2.96Å^, and SYN^P3212_3.29Å^ structures **(Figure S9A,D,G, Supporting Information)**.

**Figure 7:**
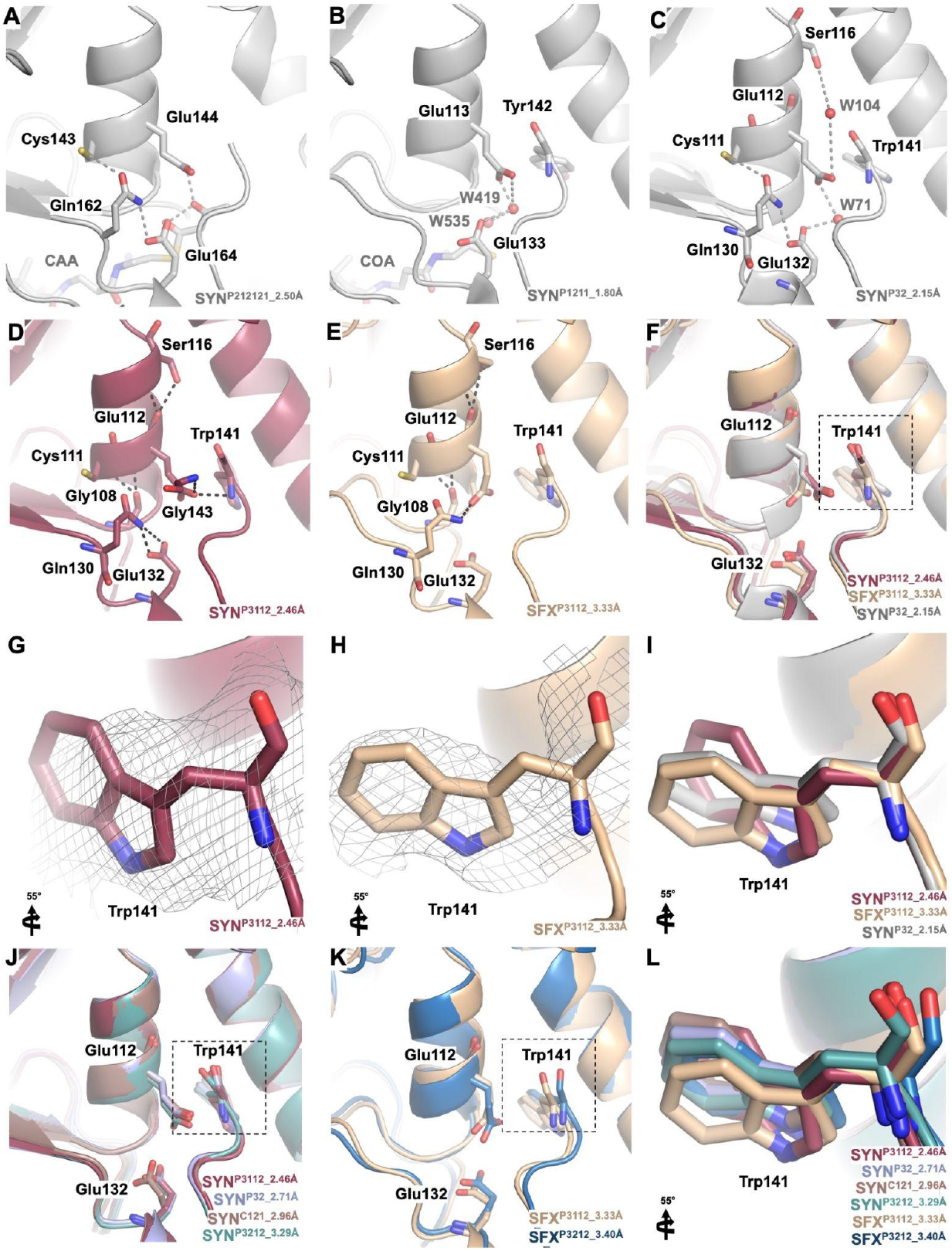
Hydrogen binding network of Nmar_1308 protein compared to previously available structures. **(A-C)** The active sites of previously available homologous structures (SYN^P212121_2.50Å^, SYN^P1211_1.80Å^ and SYN^P32_2.15Å^; PDB IDs: 1DUB, 5ZAI, and 7EUM, respectively) are represented and colored in gray. **(D-E)** The active sites of SYNP^3112_2.46Å^ and SFX^P312_3.33Å^ structures are represented and colored raspberry and wheat, respectively. Hydrogen binding interactions of Glu112 and Glu132 are indicated with dashed lines. Water molecules are shown in red spheres. **(F)** SYN^P32_2.15Å^ structure is superposed with SYN^P3112_2.46Å^ and SFX^P312_3.33Å^ structures. **(G-I,L)** Trp141 residue in the substrate binding pocket is shown in stick representation to indicate conformational changes. 2Fo-Fc electron density map is contoured at the 1σ level and colored in gray. While all cryogenic structures are superposed in panel **G**, ambient temperature structures are superposed in panel **K**.

For the ambient temperature SFX^P3112_3.33Å^ structure, only Glu112 interacts with Gly108, Ser116, and Gln130 **(Figure 7E);** and in the SFX^P3212_3.40Å^ structure, Glu112 interacts with the residues Gly108 and Ser116. **(Figure S9J, Supporting Information)**. In contrast, Glu132 does not interact with the residue Glu130 for both ambient temperature structures. These findings indicate the loss of interaction with Trp141 and Gln 130 at ambient temperature. The superposition of the SFX^P3112_3.33Å^ structure with SYN^P3112_2.46Å^ and SYN^P32_2.15Å^ structures indicated significant conformational changes on the residues Glu112, Glu132, and Trp141 **(Figure 7F)**. These conformational changes may lead to disruption or reconfiguration of the interactions.

Based on *Destan et al*., less conserved residues (Cys111, Ser116, and Gln130) have an assisting role in the bifunctionality of Nmar_1308 protein.^**[14]**^ As the Cys111-Gln130 interaction was observed in the hydratase and bifunctional structure (SYN^P212121_2.50Å^ and SYN^P32_2.15Å^; PDB IDs: 1DUB & 7EUM, respectively) during the CCAH reaction with the addition of water, this interaction disappeared in our structure with the removal of the water. On the other side, Ser116-Glu112 interaction was observed in our structures while Ser116 interacts with Glu112 through a water molecule (W104) in the SYN^P32_2.15Å^ structure.

Trp141 (Tyr142 in 3HPD of *M. sedula* (SYN^P1211_1.80Å^; PDB ID: 5ZAI)) plays a key role in limiting the size of the substrate in the binding pocket, allowing the binding of short substrates. In contrast, this residue is replaced by smaller side chain residues such as alanine (*Rattus norvegicus* (SYN^P212121_2.50Å^; PDB ID: 1DUB)) to allow longer chain substrates to enter **(Figure 8)**. Significantly altered conformations were observed for the residue Trp141 in both cryogenic and ambient temperature structures **(Figure 6G-L; Figure S9, S10, Supporting Information)**. These structures offer snapshots from the different conformational states of a bifunctional Nmar_1308 protein, revealing the missing details of enzyme function.

**Figure 8:**
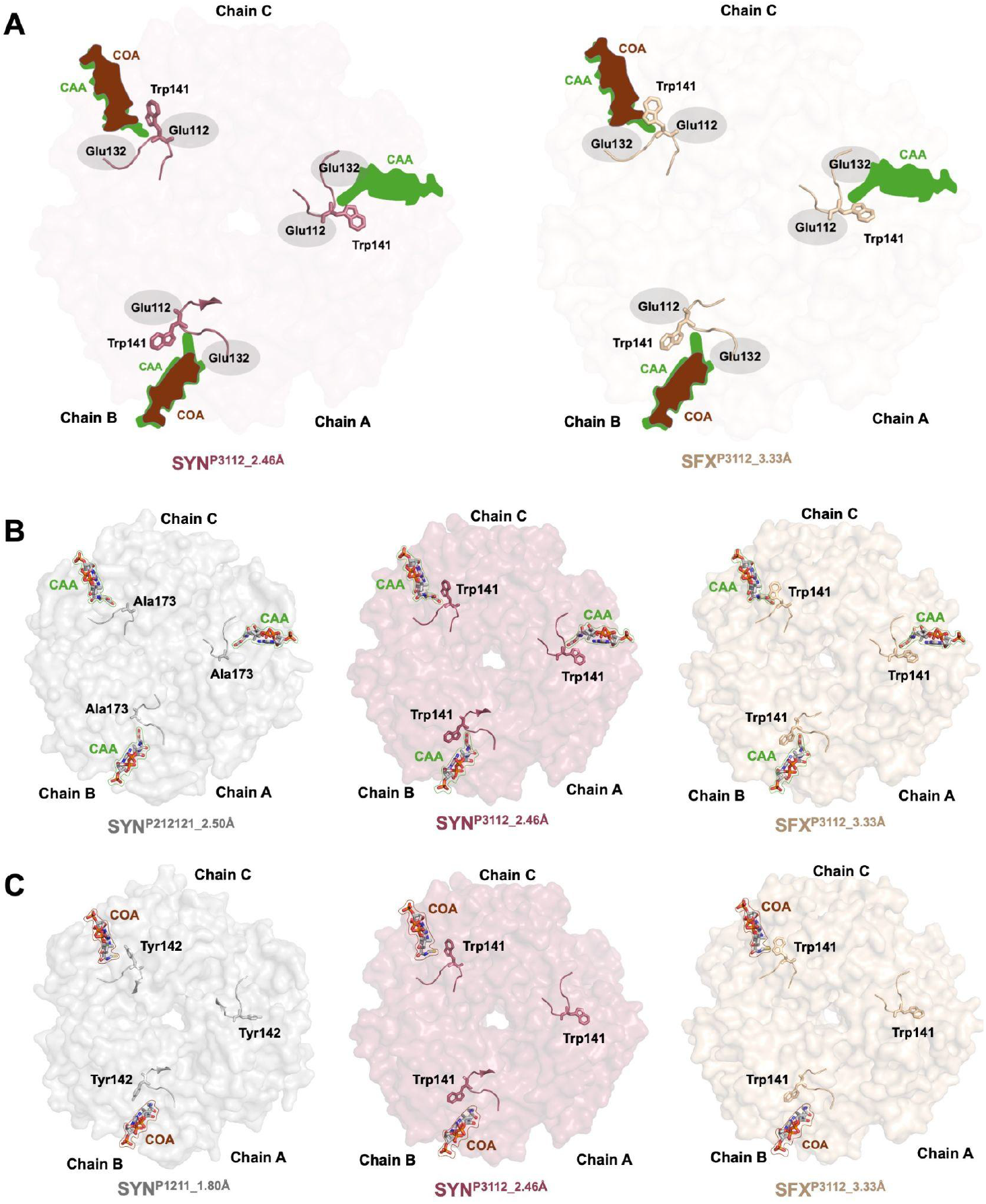
Representation of substrate entrance to the active site of the Nmar_1308 protein. **(A)** The critical position of the residue, Trp141, which limits the substrate size at the active site is shown for both cryogenic temperature SYN^P3112_2.46Å^ and ambient temperature SFX^P3112_3.33Å^ structures. **(B)** The bulky residue, Trp141, is replaced by Ala173 in the presence of a long substrate such as acetoacetyl-coenzyme A (CAA) (PDB ID: 1DUB). **(C)** For the short cofactor such as coenzyme A (COA) (PDB ID: 5ZAI), Trp141 is replaced by another bulky residue, Tyr142.

## Conclusions

The data presented herein provide new insight into the mechanism of action of a bifunctional enzyme from *N. maritimus*. The Nmar_1308 protein plays a critical role in the 3HP/4HB cycle, which is recognized as one of the most efficient CO_2_ fixation pathways, through its dual functionality as both a CCAH and a 3HPD. The obtained structures during this study reveal previously unobserved conformational changes at the active site in both cryogenic and ambient temperatures. While the previous cryogenic temperature structure (SYN^P32_2.15Å^; PDB ID: 7EUM) highlights the importance of the critical water molecule (W71) during the hydratase reaction, our structures capture the structural dynamics of the dehydratase reaction, revealing the missing details of the enzyme’s mechanism of action. Additionally, a previously disordered region, including the critical residue Ala64, was observed in a more stable conformation, providing valuable insights into the structural dynamics of the active site and the mechanism of substrate entry. The comprehensive structural analysis of the Nmar_1308 protein at both cryogenic and, for the first time, ambient temperatures offers a foundational framework for future biotechnological applications and enhances our understanding of archaeal carbon fixation.

## Materials and Methods

### Protein sample preparation

The codon-optimized Nmar_1308 gene construct which is previously described in *Destan et al*. was used for obtaining the Nmar_1308 protein.^**[14]**^ The plasmid was transformed into *Escherichia coli*, Rosetta2™ BL21 strain, and the transformed cells were grown overnight on agar plates containing 30 µg/ml chloramphenicol and 50 µg/ml kanamycin at 37°C. For large-scale protein purification, overnight cultures were initiated from single colonies on agar plates, grown in LB media, and then diluted 1:100 into 2L cultures. When the cultures reached an optical density of 0.8 at 600 nm, IPTG was added to a final concentration of 0.7 mM to induce protein expression and incubated overnight at 16 °C. Cells were harvested by centrifuge at 3500 rpm. Then, cell pellets were resuspended in a lysis buffer (250 mM NACI, 50 mM Tris pH 7.5, 0.01% Triton X-100) and sonicated (Branson W250 sonifier, USA). After the sonication step, cell lysate was centrifuged at 35000 rpm for 1 hr at 4°C. Protein purification was performed using Ni-NTA affinity resin (GE Healthcare). The column was initially washed with a binding buffer (150 mM NaCl, 20 mM TRIS pH 7.5), and the bound protein was eluted with an elution buffer (150 mM NaCl, and 20 mM TRIS pH 7.5, 250 mM Imidazole). Thrombin was added to cleave the N-terminal histidine tag, and reverse Ni-NTA chromatography was used to remove the cleaved tag. The enzyme-containing fractions, free of the affinity tag, were pooled, concentrated, and subjected to final purification using a Superdex 200 (S200) column. The final product was concentrated to 10 mg/ml before crystallization and protein concentration was measured by UV absorption at 280 nm using a Nanodrop spectrophotometer. The purity of the protein was assessed by SDS-PAGE.

### Crystallization

Crystallization was performed by using the sitting-drop vapor diffusion micro-batch under the oil method. The purified Nmar_1308 protein at 10 mg/ml was mixed with ∼3500 crystallization conditions with a 1:1 volumetric ratio for screening, in the 72-well Terasaki^TM^ plates (Greiner-Bio, Germany). The best crystals were obtained by using Morpheus I and Helix I commercial crystallization kits (Molecular Dimensions, UK) **(Figure S1; Table S1, Supporting Information)**. For SFX experiments, the batch crystallization volume was scaled up to a total of 1.5 ml.

### Cryogenic-temperature data collection and processing at SPring-8

Cryogenic data was collected at the BL32XU beamline at SPring-8 using the automated data-collection system ZOO described by *Hirata et al*.^**[17]**^ Before data collection, crystals were frozen in liquid nitrogen and pucks were replaced into the dewar which is filled with liquid nitrogen in the hutch. All data sets were acquired using a continuous helical scheme with 360° of oscillation and the following experimental parameters: oscillation width, 0.1°; exposure time, 0.02 s; beam size, 10 µm (horizontal) × 15 µm (vertical); wavelength, 1 Å; average dose per crystal volume, 10 MGy; detector, EIGER X 9M (Dectris); temperature, 100 K. The data obtained were automatically processed by XDS^**[18]**^ in KAMO^**[17]**^.

### SFX data collection and processing at SACLA

Serial femtosecond X-ray diffraction data at ambient temperature (298 K) were collected at the Experimental Hutch 3 of SACLA Beamline 2 using 10 keV X-rays with a pulse duration of <10 fs and a repetition rate of 30 Hz. The focused beam size is 1.5 µm-to-diameter. Before data collection, crystals were crushed to obtain microcrystals. Then, the crystal slurry was centrifuged at 3000 rpm for 5 min to reach 10^8^ crystal density as described in *Destan et al*.^**[19]**^ The crystals were mixed with grease and transferred into a 200 μL sample cartridge carefully by using a spatula. The prepared cartridge was loaded into the high-viscosity cartridge-type (HVC) injector with a nozzle of 75 μm (inner diameter), and the HVC injector was replaced into the He-gas-filled diffraction chamber of the DAPHNIS instrument.^**[20,21]**^ Diffraction patterns were recorded by MPCCD detector (50 mm distance) and the sample slurry (mixed with grease) was injected at a flow rate of 0.79 µL min−1. Diffraction images were processed by using Cheetah^**[22]**^, which was modified for processing SFX data at SACLA^**[23]**^, and CrystFEL^**[24]**^ during data collection. For the SFX^P312_3.33Å^ structure, four injections were done and a total of 31646 out of 109336 hits were indexed. For the SFX^P312_3.40Å^ structure, five injections were done and a total of 33994 out of 112464 hits were indexed.

### Structure determination and refinement

The four synchrotron structures in space groups P3112, P32, C121, and P3212; and two SFX structures in space groups P3112, and P3212 were determined by using the molecular replacement program *PHASER*^**[25]**^ implemented in the *PHENIX*^**[26]**^ suite. The coordinates of the previously published apo structure of Nmar 1308 at cryogenic temperature (PDB ID: 7EUM) were used for initial rigid body refinement. After the simulated annealing refinement, TLS parameters and individual atomic coordinates were further refined. Additionally, composite omit map refinement was carried out using *PHENIX* to detect possible locations of the altered side chains and water molecules, which were examined manually in *COOT*^**[27]**^. The data collection and structure refinement statistics are summarized in Table 1.

## Acknowledgments

This publication has been produced benefiting from the 2244 Industrial PhD Fellowship Program Funding Program of the Scientific and Technological Research Council of Turkey (TÜBİTAK) (Project Numbers: 119C132). However, the entire responsibility of the publication belongs to the owner of the publication. The financial support received from TÜBİTAK does not mean that the content of the publication is approved in a scientific sense by TÜBİTAK. Portions of this research were carried out at the SPring-8 Angstrom Compact free electron LAser, Japan (SACLA) and supported by the SACLA Research Support Program for Graduate Students (Proposal numbers: 2023B8058 and 2023B2761).

## Author Contributions

The project was initiated and coordinated by S.W., H.D.; Protein sample and diffracting crystals were obtained by E.D.; Crystallography applications at *SPring-8/SACLA* were supported by E.D., J.K., T.T. and M.Y; and in beamtime, samples were prepared by E.D.; SFX data at *SACLA* was collected by E.D. and J.K.; Synchrotron data at *SPring-8* was collected by E.D., J.K., and T.T with the help of Y.K.; The data processing was performed for SFX structures by E.D. and for synchrotron structures by H.M.; Structures were determined and refined by E.D.; Data were interpreted and the manuscript was written by E.D., I.Y., B.B.T., C.K., Z.N., S.D., Y.Y., C.A.F., S.W., and H.D.; All of the authors read and acknowledged the manuscript.

## Data availability

Coordinates and structure factors have been deposited in the RCSB Protein Data Bank under accession codes 9UL8, 9UL9, 9ULA and 9ULB, which correspond to the cryogenic temperature structures (SYN^P3112_2.46Å^, SYN^P32_2.71Å^, SYN^C121_2.96Å^, and SYN^P3212_3.29Å^, respectively); and 9ULC and 9ULD, which correspond to the ambient structures (SFX^P3112_3.33^ Å and SFX^P3212_3.40Å^, respectively). Any remaining information can be obtained from the corresponding author upon request.

## Supporting Information

Figures S1-10; Table S1.

## References

[1] M. Könneke, D. M. Schubert, P. C. Brown, M. Hügler, S. Standfest, T. Schwander, L. Schada von Borzyskowski, T. J. Erb, D. A. Stahl, I. A. Berg, Proc. Natl. Acad. Sci. 2014, 111, 8239.

[2] A. E. Ingalls, S. R. Shah, R. L. Hansman, L. I. Aluwihare, G. M. Santos, E. R. M. Druffel, A. Pearson, Proc. Natl. Acad. Sci. U. S. A. 2006, 103, 6442.

[3] Q. Yin, K. He, G. Collins, J. De Vrieze, G. Wu, npj Clean Water 2024, 7, 52.

[4] P. Friedlingstein, M. Meinshausen, V. K. Arora, C. D. Jones, A. Anav, S. K. Liddicoat, R. Knutti, J. Clim. 2014, 27, 511.

[5] C. B. Walker, J. R. de la Torre, M. G. Klotz, H. Urakawa, N. Pinel, D. J. Arp, C. Brochier-Armanet, P. S. G. Chain, P. P. Chan, A. Gollabgir, J. Hemp, M. Hügler, E. A. Karr, M. Könneke, M. Shin, T. J. Lawton, T. Lowe, W. Martens-Habbena, L. A. Sayavedra-Soto, D. Lang, S. M. Sievert, A. C. Rosenzweig, G. Manning, D. A. Stahl, Proc. Natl. Acad. Sci. 2010, 107, 8818.

[6] A. E. Santoro, C. L. Dupont, R. A. Richter, M. T. Craig, P. Carini, M. R. McIlvin, Y. Yang, W. D. Orsi, D. M. Moran, M. A. Saito, Proc. Natl. Acad. Sci. 2015, 112, 1173.

[7] A. H. Treusch, S. Leininger, A. Kletzin, S. C. Schuster, H. Klenk, C. Schleper, Environ. Microbiol. 2005, 7, 1985.

[8] W. Martens-Habbena, P. M. Berube, H. Urakawa, J. R. de la Torre, D. A. Stahl, Nature 2009, 461, 976.

[9] B. Kraft, N. Jehmlich, M. Larsen, L. A. Bristow, M. Könneke, B. Thamdrup, D. E. Canfield, Science (80-. ). 2022, 375, 97.

[10] J. Johnson, B. B. Tolar, B. Tosun, Y. Yoshikuni, C. A. Francis, S. Wakatsuki, H. DeMirci, Commun. Biol. 2024, 7, 1364.

[11] I. A. Berg, Appl. Environ. Microbiol. 2011, 77, 1925.

[12] L. Liu, D. M. Schubert, M. Könneke, I. A. Berg, Front. Microbiol. 2021, 12.

[13] D. Lee, K.-J. Kim, Sci. Rep. 2018, 8, 10692.

[14] E. Destan, B. Yuksel, B. B. Tolar, E. Ayan, S. Deutsch, Y. Yoshikuni, S. Wakatsuki, C. A. Francis, H. DeMirci, Sci. Rep. 2021, 11, 22849.

[15] I. A. Berg, D. Kockelkorn, W. H. Ramos-Vera, R. F. Say, J. Zarzycki, M. Hügler, B. E. Alber, G. Fuchs, Nat. Rev. Microbiol. 2010, 8, 447.

[16] J. K. Kim, H. Jeong, J. Seo, K. H. Kim, D. Min, C. U. Kim, Acta Crystallogr. Sect. D, Struct. Biol. 2024, 80, 686.

[17] K. Hirata, K. Yamashita, G. Ueno, Y. Kawano, K. Hasegawa, T. Kumasaka, M. Yamamoto, Acta Crystallogr. Sect. D, Struct. Biol. 2019, 75, 138.

[18] W. Kabsch, B. A. T., D. K., K. P. A., D. K., M. S., R. R. B. G., E. P., F. S., W. K., K. W., K. W., K. W., K. W., K. P., W. M. S., Acta Crystallogr. Sect. D Biol. Crystallogr. 2010, 66, 125.

[19] E. Destan, E. Turkut, A. Aldeniz, J. Kang, T. Tosha, M. Yabashi, B. Yilmaz, A. C. Timucin, H. Matsuura, Y. Kawano, I. Cinkaya, H. DeMirci, Small Struct. 2025, 2400680.

[20] K. Tono, E. Nango, M. Sugahara, C. Song, J. Park, T. Tanaka, R. Tanaka, Y. Joti, T. Kameshima, S. Ono, T. Hatsui, E. Mizohata, M. Suzuki, T. Shimamura, Y. Tanaka, S. Iwata, M. Yabashi, J. Synchrotron Radiat. 2015, 22, 532.

[21] Y. Shimazu, K. Tono, T. Tanaka, Y. Yamanaka, T. Nakane, C. Mori, K. Terakado Kimura, T. Fujiwara, M. Sugahara, R. Tanaka, R. B. Doak, T. Shimamura, S. Iwata, E. Nango, M. Yabashi, J. Appl. Crystallogr. 2019, 52, 1280.

[22] A. Barty, R. A. Kirian, F. R. N. C. Maia, M. Hantke, C. H. Yoon, T. A. White, H. Chapman, J. Appl. Crystallogr. 2014, 47, 1118.

[23] T. Nakane, Y. Joti, K. Tono, M. Yabashi, E. Nango, S. Iwata, R. Ishitani, O. Nureki, J. Appl. Crystallogr. 2016, 49, 1035.

[24] T. A. White, R. A. Kirian, A. V. Martin, A. Aquila, K. Nass, A. Barty, H. N. Chapman, J. Appl. Crystallogr. 2012, 45, 335.

[25] A. J. McCoy, R. W. Grosse-Kunstleve, P. D. Adams, M. D. Winn, L. C. Storoni, R. J. Read, J. Appl. Crystallogr. 2007, 40, 658.

[26] P. D. Adams, P. V. Afonine, G. Bunkóczi, V. B. Chen, I. W. Davis, N. Echols, J. J. Headd, L.-W. Hung, G. J. Kapral, R. W. Grosse-Kunstleve, A. J. McCoy, N. W. Moriarty, R. Oeffner, R. J. Read, D. C. Richardson, J. S. Richardson, T. C. Terwilliger, P. H. Zwart, Acta Crystallogr. Sect. D Biol. Crystallogr. 2010, 66, 213.

[27] P. Emsley, K. Cowtan, Acta Crystallogr. Sect. D Biol. Crystallogr. 2004, 60, 2126.

