## Supporting Information for "Deciphering the altered conformational states of bifunctional thaumarchaeal crotonyl-CoA hydratase and 3-hydroxypropionyl-CoA dehydratase from *Nitrosopumilus maritimus*"

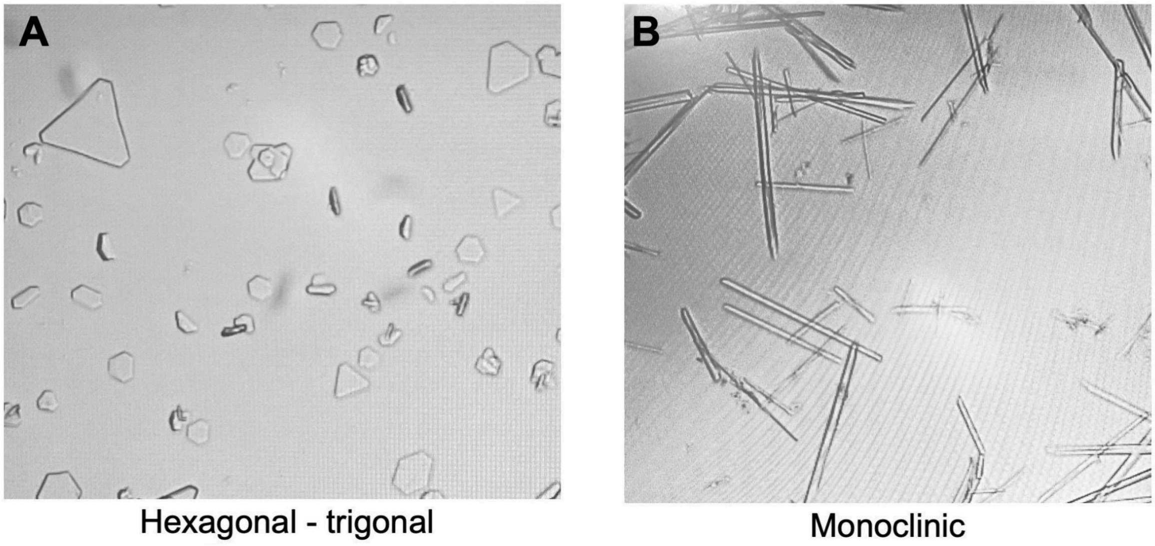


**Figure S1:** Crystal images of Nmar_1308 protein with two lattice types. The crystal structures with P3112, P31, and P312 space groups are obtained from crystals in panel **A** while the crystal structure with C121 space group is obtained from the crystals in panel **B**.

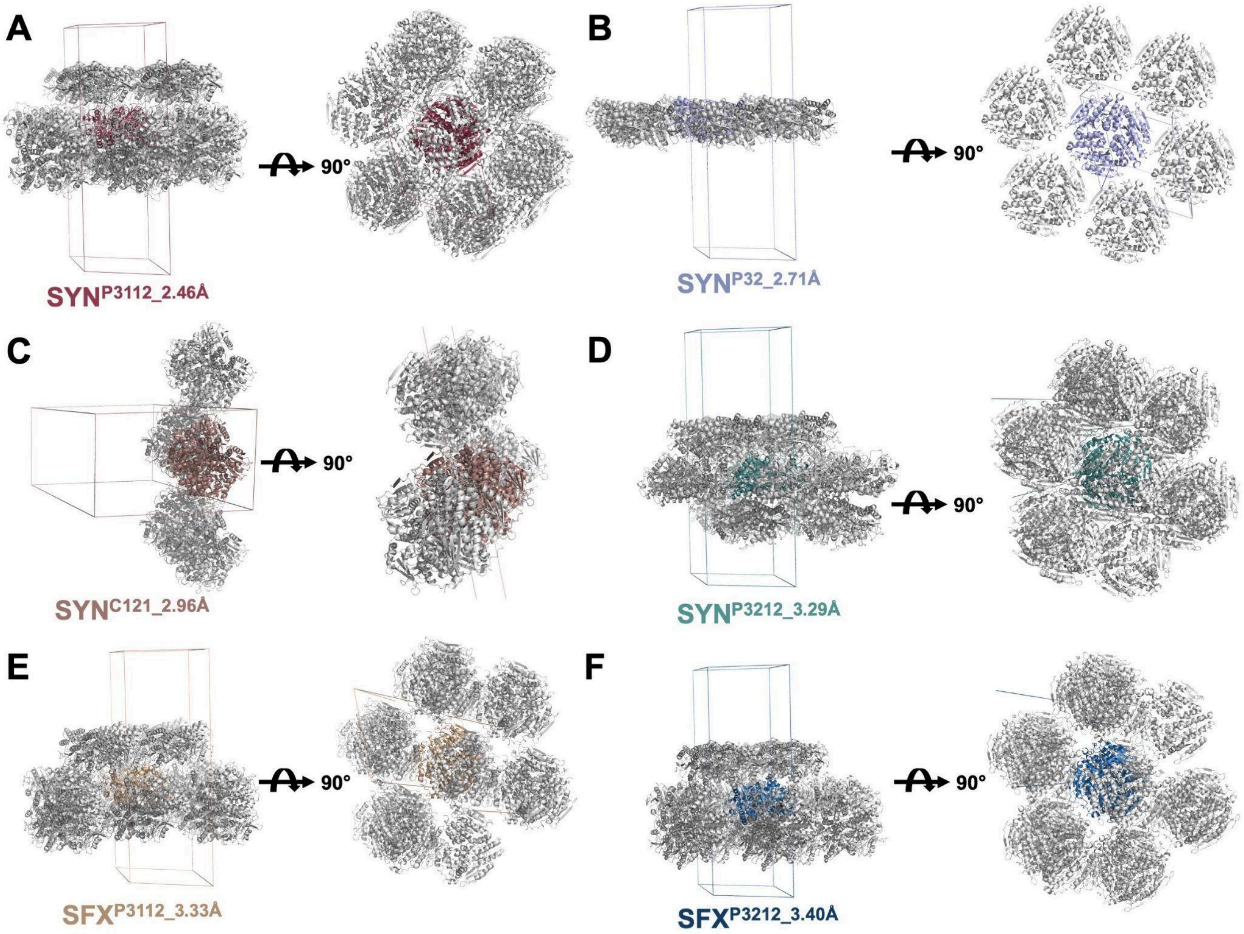


**Figure S2:** Representation of crystal symmetries around 20 Å.

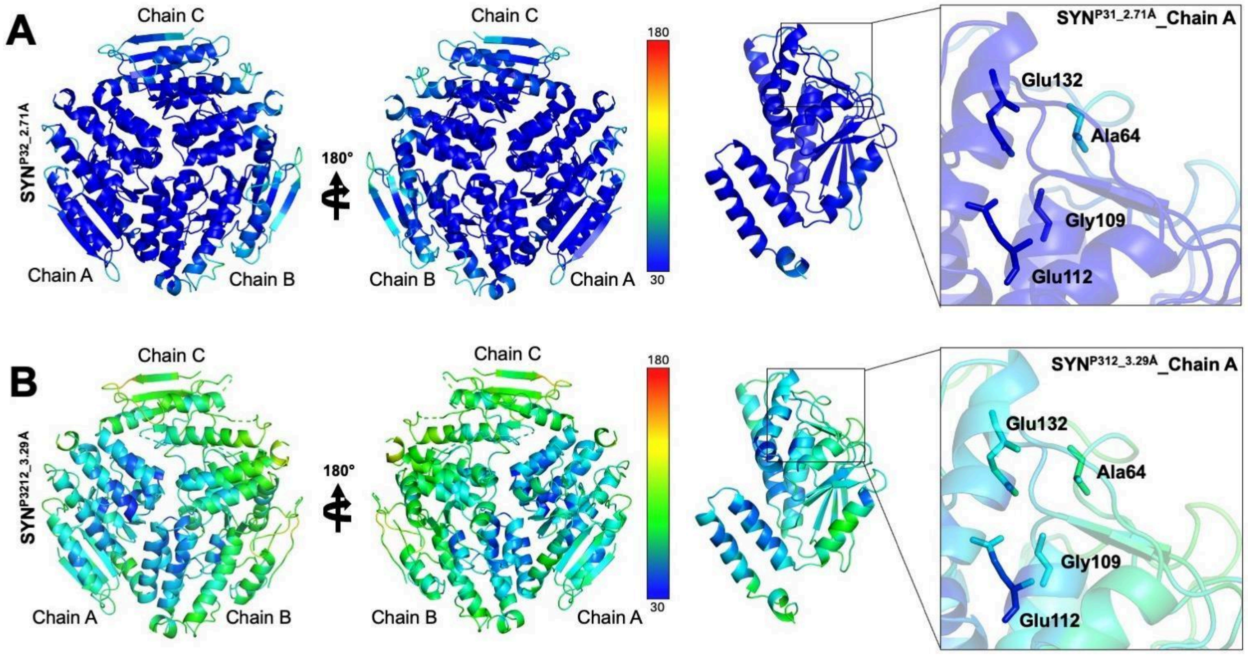


**Figure S3:** Apo-form Nmar_1308 protein crystal structures based on b-factor. **(A-B)** All structures are colored based on the b-factor (Spectrum range: 30 to 180). Chain A of each structure is represented to compare the stability around the substrate binding pocket.

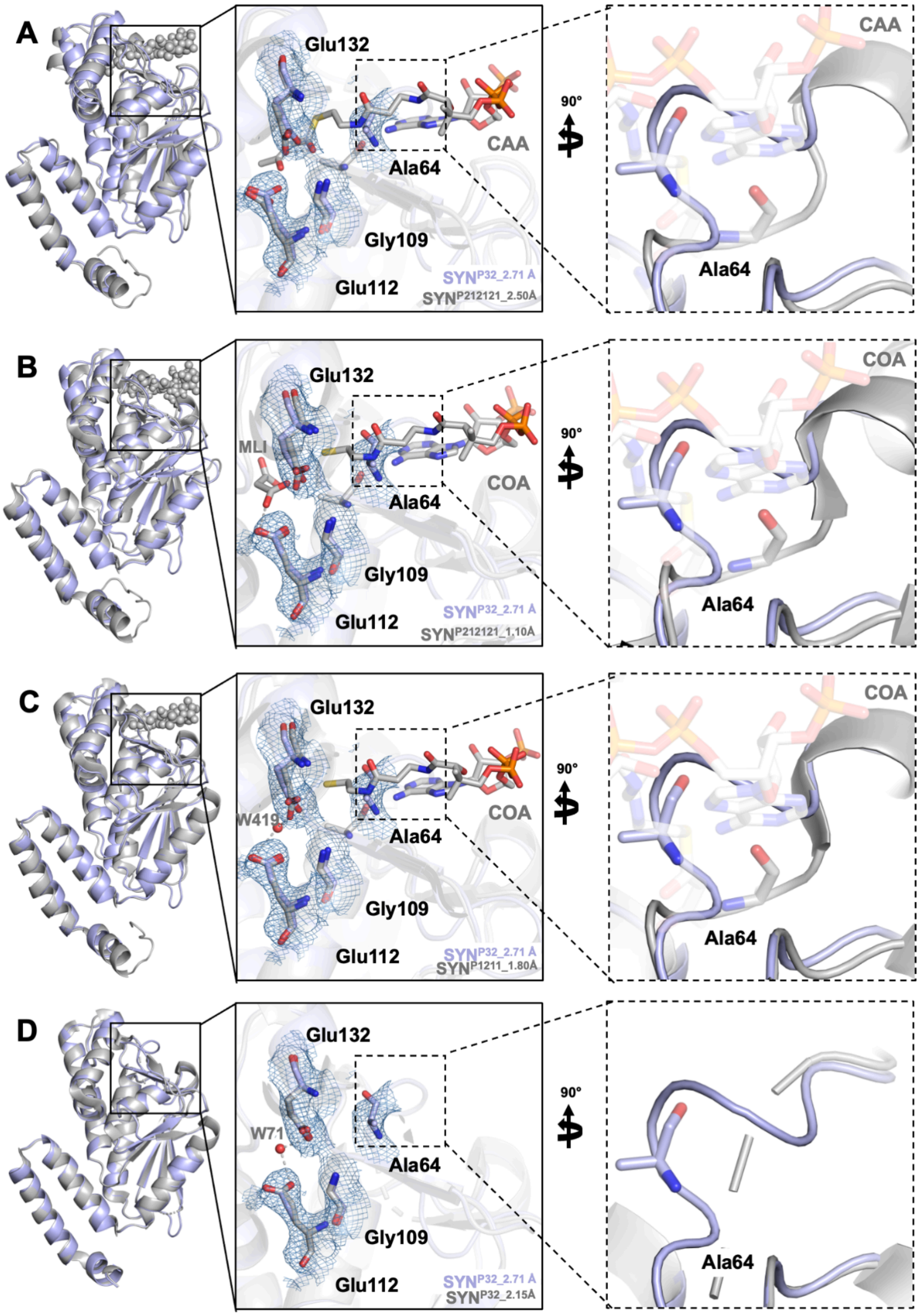


**Figure S4:** Comparison of SYN$^{P32_2.71Å}$ structure with previously available homologous hydratase/dehydratase structures. **(A-D)** To compare the substrate binding pocket, the 2.71 Å synchrotron structure is superimposed with a hydratase structure (SYN$^{P212121_2.50Å}$; PDB ID: 1DUB), dehydratase structure (SYN$^{P212121_1.10Å}$; PDB ID: 5JBX), and the biofunctional enzyme from *Metallosphaera sedula* (SYN$^{P1211_1.80Å}$; PDB ID: 5ZAI) and *Nitrosopumilus maritimus* (SYN$^{P32_2.15Å}$; PDB ID: 7EUM). 2Fo-Fc electron density map for the active site residues in the SYN$^{P32_2.71Å}$ structure is contoured at the 1σ level and colored in skyblue.

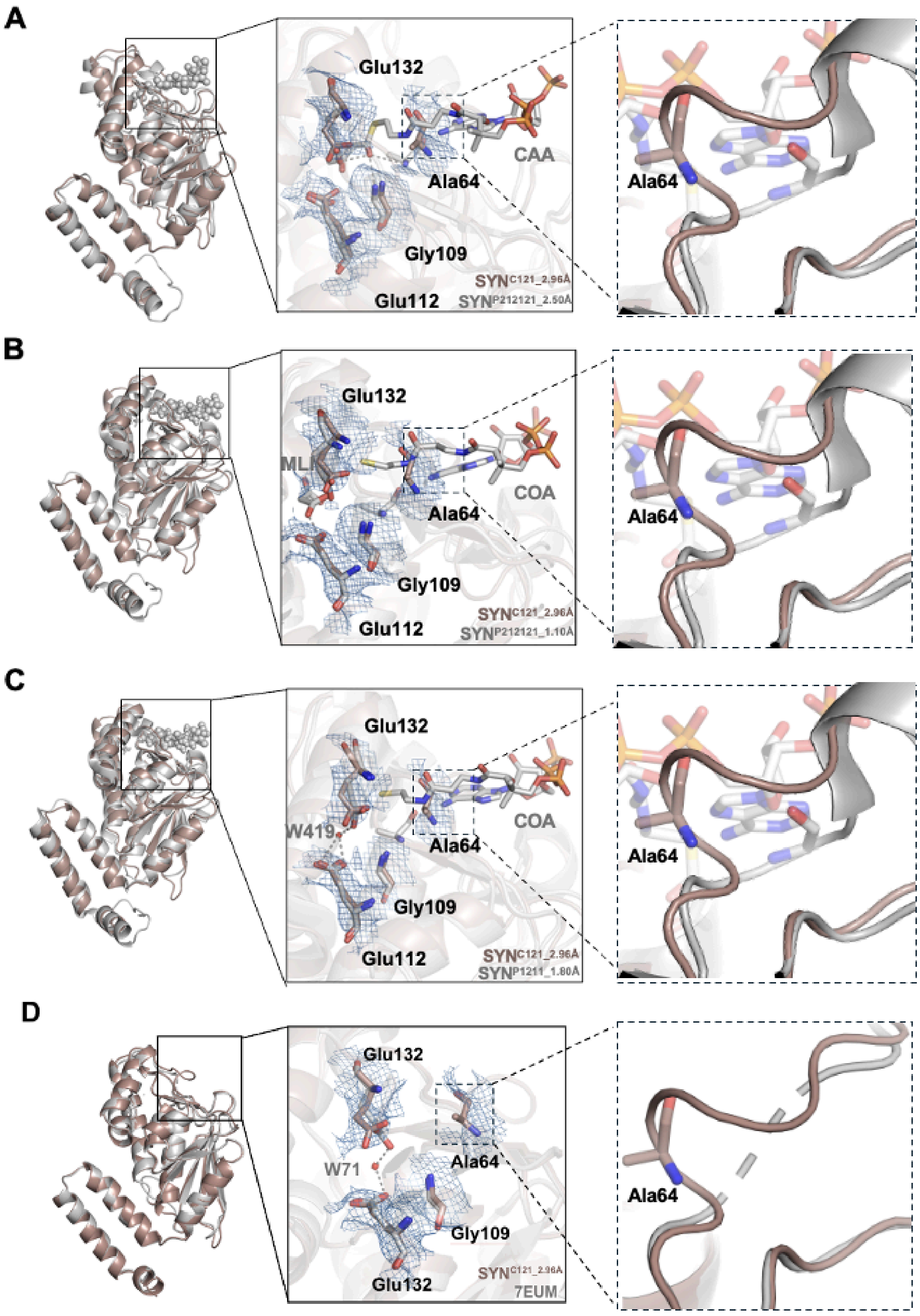


**Figure S5:** Comparison of SYN$^{\text{C121_2.96Å}}$ structure with previously available homologous hydratase/dehydratase structures. **(A-D)** To compare the substrate binding pocket, the 2.96 Å synchrotron structure is superimposed with a hydrates structure (SYN$^{\text{P212121_2.50Å}}$; PDB ID: 1DUB), dehydrates structure (SYN$^{\text{P212121_1.10Å}}$; PDB ID: 5JBX), and biofunctional enzyme from *Metallosphaera sedula* (SYN$^{\text{P1211_1.80Å}}$; PDB ID: 5ZAI) and *Nitrosopumilus maritimus* (SYN$^{\text{P32_2.15Å}}$; PDB ID: 7EUM). 2Fo-Fc electron density map for the active site residues in the SYN$^{\text{C121_2.96Å}}$ structure is contoured at the 1σ level and colored in skyblue.

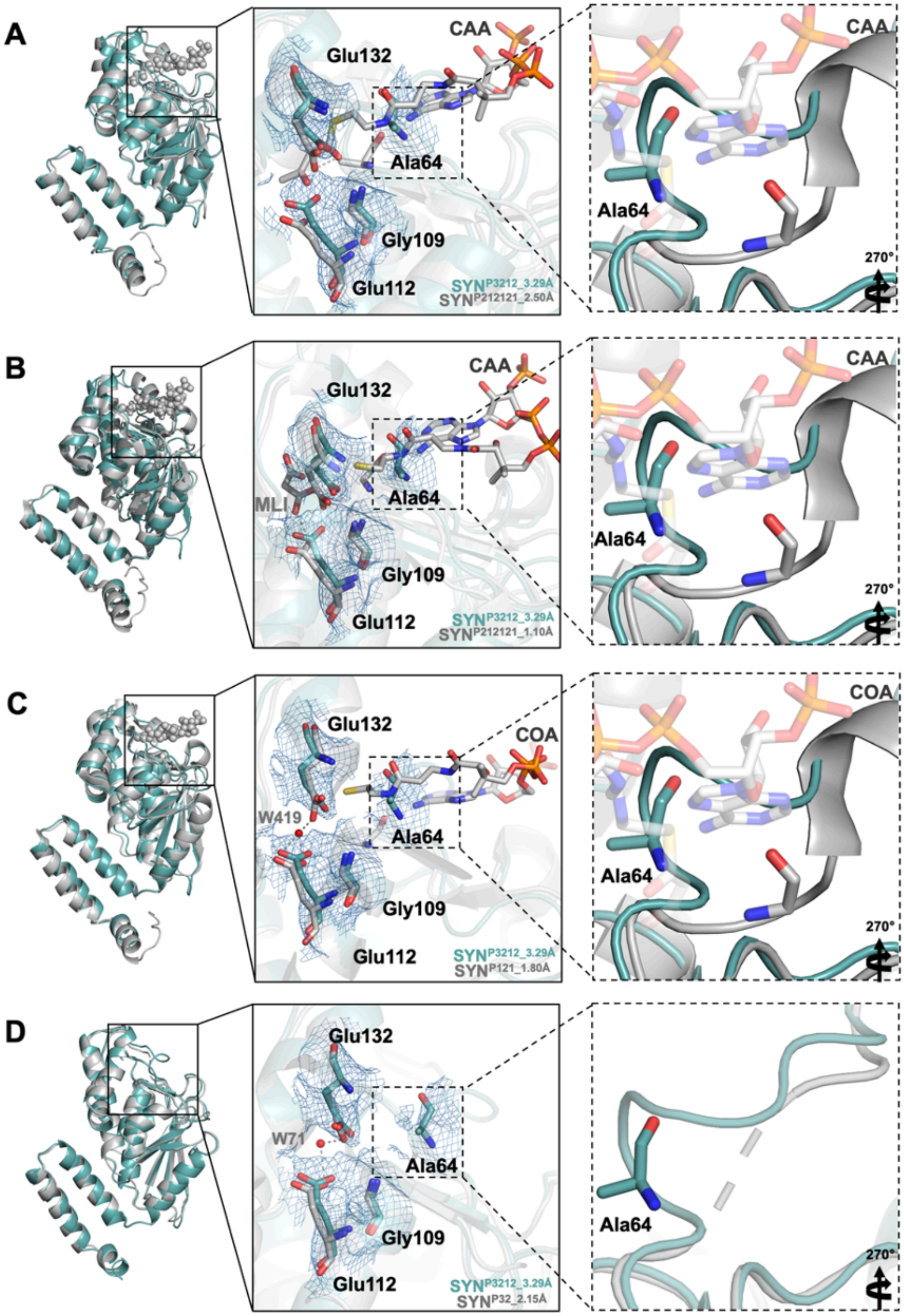


**Figure S6:** Comparison of SYN$^{P3212_3.29Å}$ structure with previously available homologous hydratase/dehydratase structures. **(A-D)** To compare the substrate binding pocket, the 3.29 Å synchrotron structure is superimposed with a hydrates structure (SYN$^{P212121_2.50Å}$; PDB ID: 1DUB), dehydrates structure (SYN$^{P212121_1.10Å}$; PDB ID: 5JBX), and biofunctional enzyme from *Metallosphaera sedula* (SYN$^{P1211_1.80Å}$; PDB ID: 5ZAI) and *Nitrosopumilus maritimus* (SYN$^{P32_2.15Å}$; PDB ID: 7EUM). 2Fo-Fc electron density map for active site residues in the SYN$^{P3212_3.29Å}$ structure is contoured at the 1σ level and colored in skyblue.

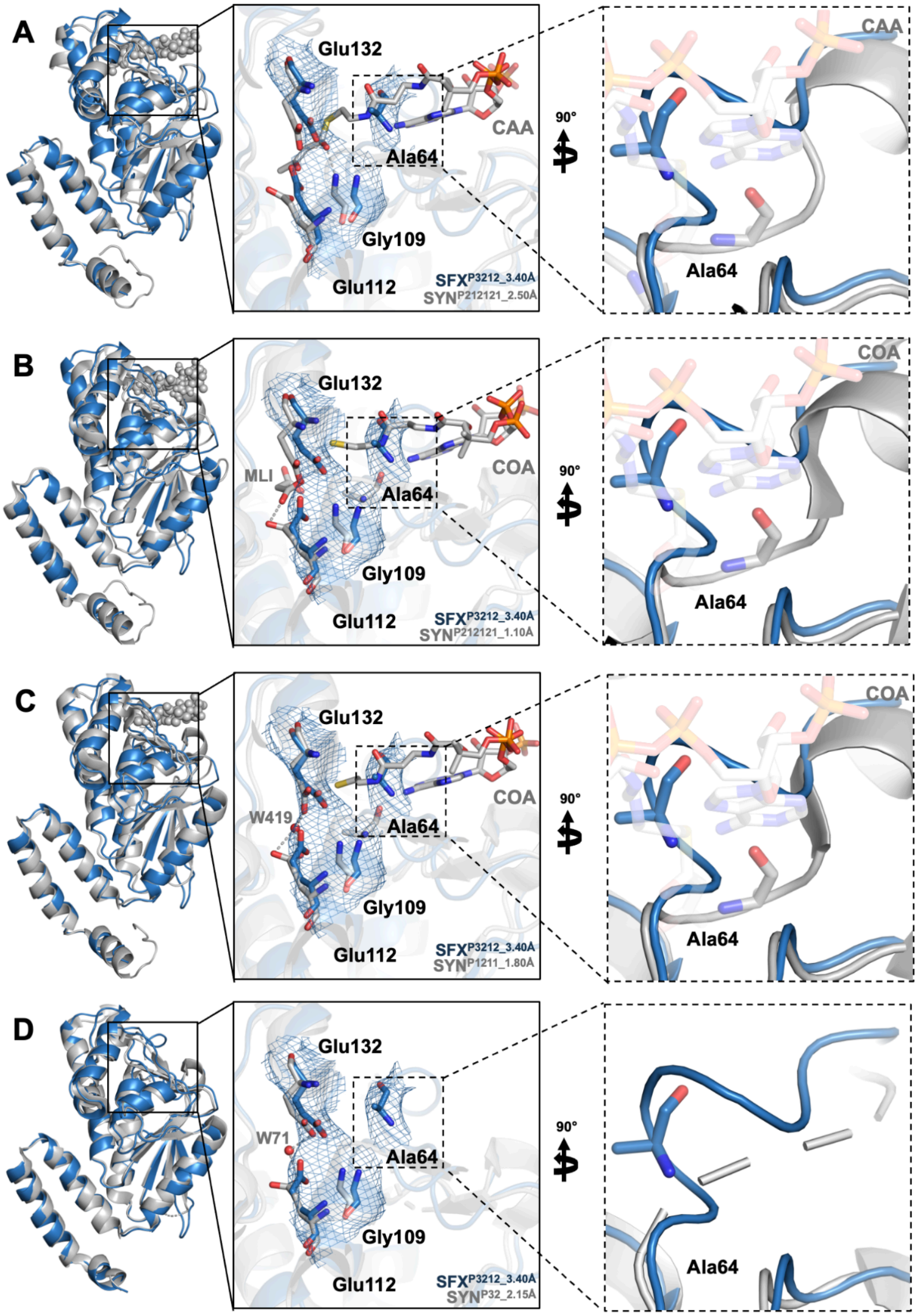


**Figure S7:** Comparison of SFX$^{P3212_3.40Å}$ structure with previously available homologous hydratase/dehydratase structures. **(A-D)** To compare the substrate binding pocket, the 3.40 Å synchrotron structure is superimposed with a hydrates structure (SYN$^{P212121_2.50Å}$; PDB ID: 1DUB), dehydrates structure (SYN$^{P212121_1.10Å}$; PDB ID: 5JBX), and biofunctional enzyme from *Metallosphaera sedula* (SYN$^{P1211_1.80Å}$; PDB ID: 5ZAI) and *Nitrosopumilus maritimus* (SYN$^{P32_2.15Å}$; PDB ID: 7EUM). 2Fo-Fc electron density map for active site residues in the SFX$^{P3212_3.40Å}$ structure is contoured at the 1σ level and colored in skyblue.

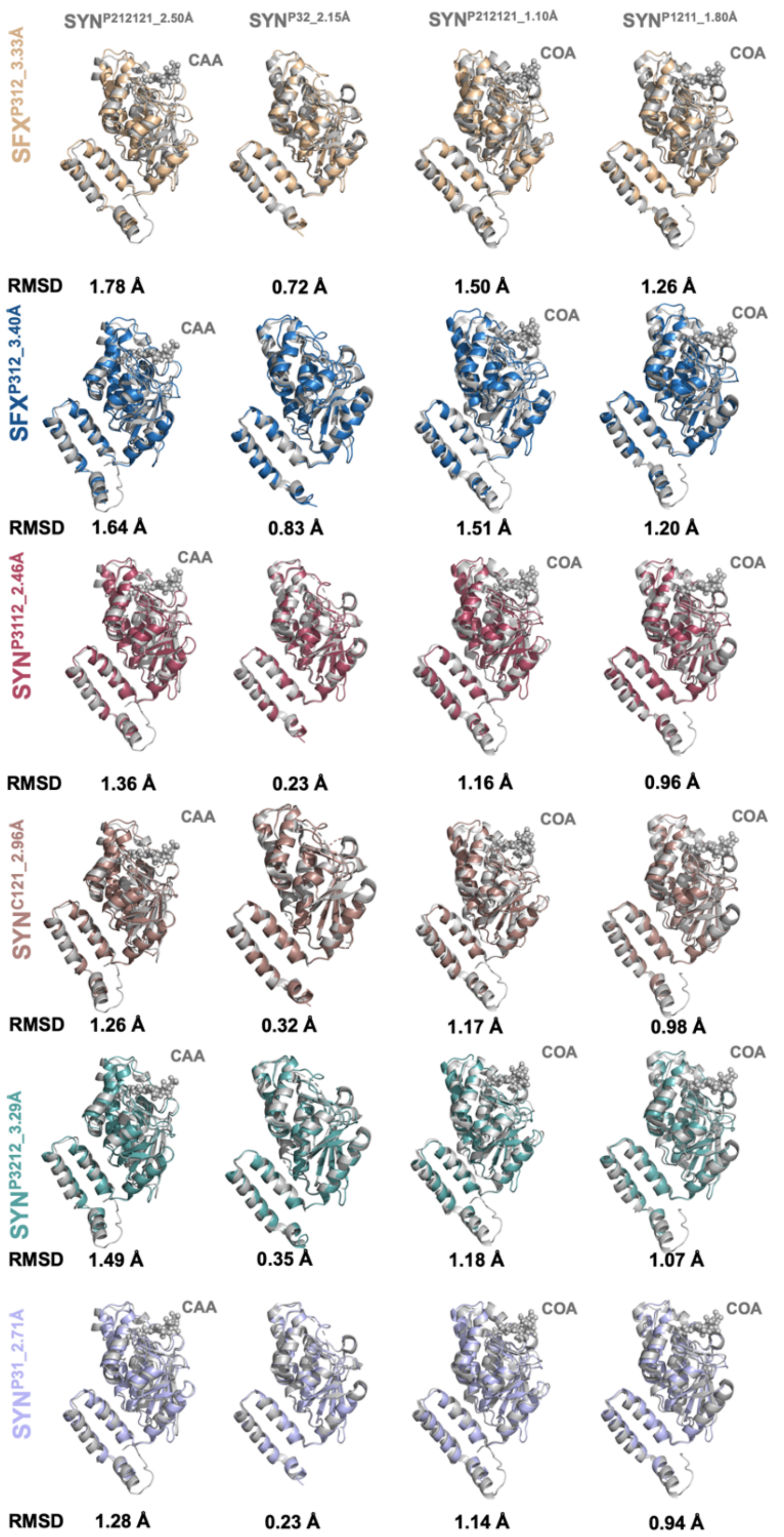


**Figure S8:** Superimposition of cryogenic and ambient temperature structures with previously available hydratase/dehydratase homologous structures. Chain A of each structure was used during superposition.

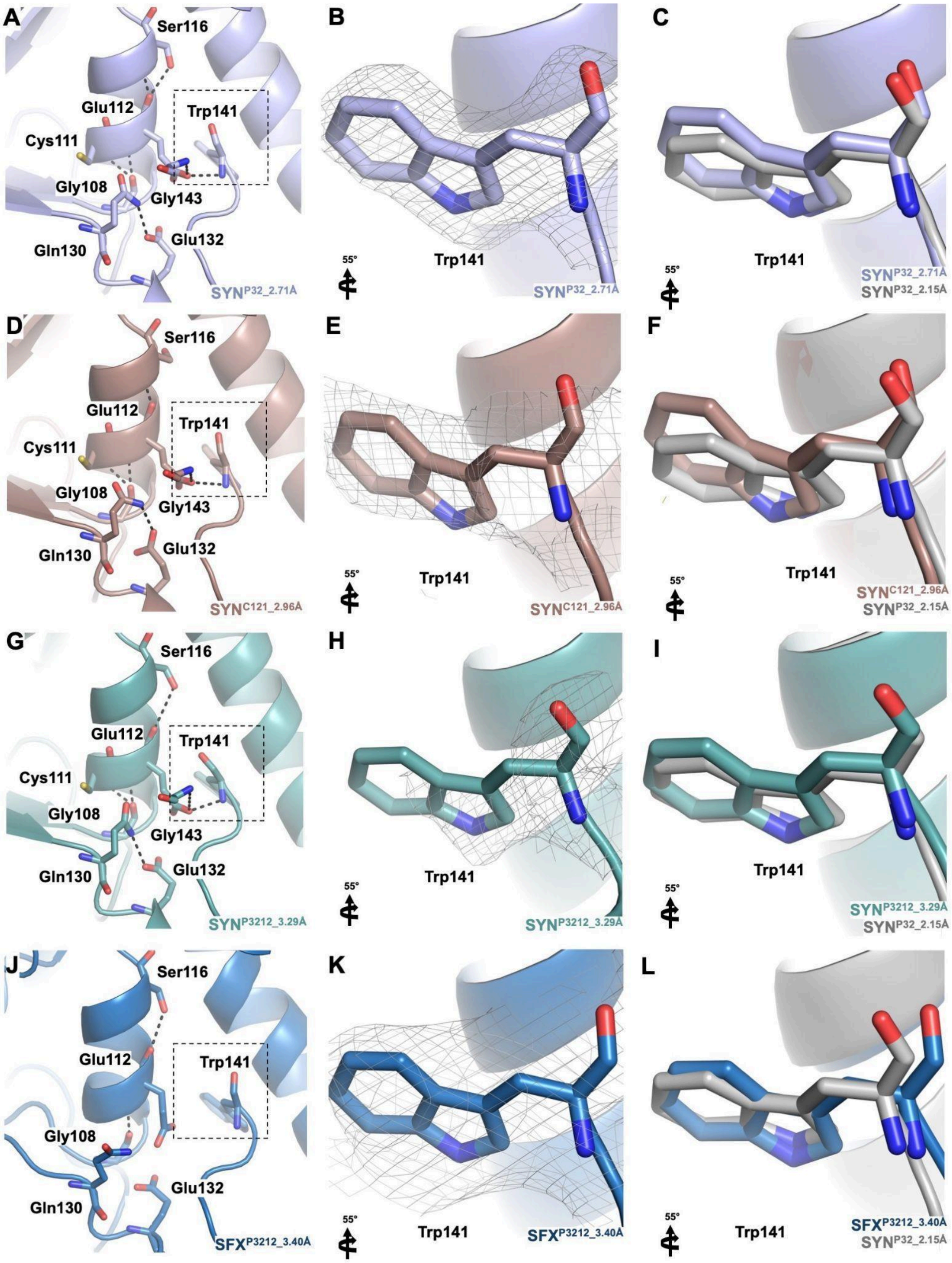


**Figure S9:** Hydrogen bonding network of Nmar_1308 protein compared to previously available structures. **(A-C)** The active site of $SYN^{P31\_2.71Å}$ and Trp141, a residue important for limiting substrate size, is represented in the substrate binding pocket. **(D-F)** The active site of $SYN^{C121\_2.96Å}$ and Trp141, a residue important for limiting substrate size, is represented in the substrate binding pocket. **(G-I)** The active site of $SYN^{P312\_3.29Å}$ and Trp141, a residue important for limiting substrate size, is represented in the substrate binding pocket. **(J-L)** The active site of $SFX^{P312\_3.40Å}$ and Trp141, a residue important for limiting substrate size, is represented in the substrate binding pocket. The previously available crystal structure of Nmar_1308 protein ($SYN^{P32\_2.15Å}$; PDB ID: 7EUM) is colored gray. 2Fo-Fc electron density map is contoured at the 1σ level and colored in gray.

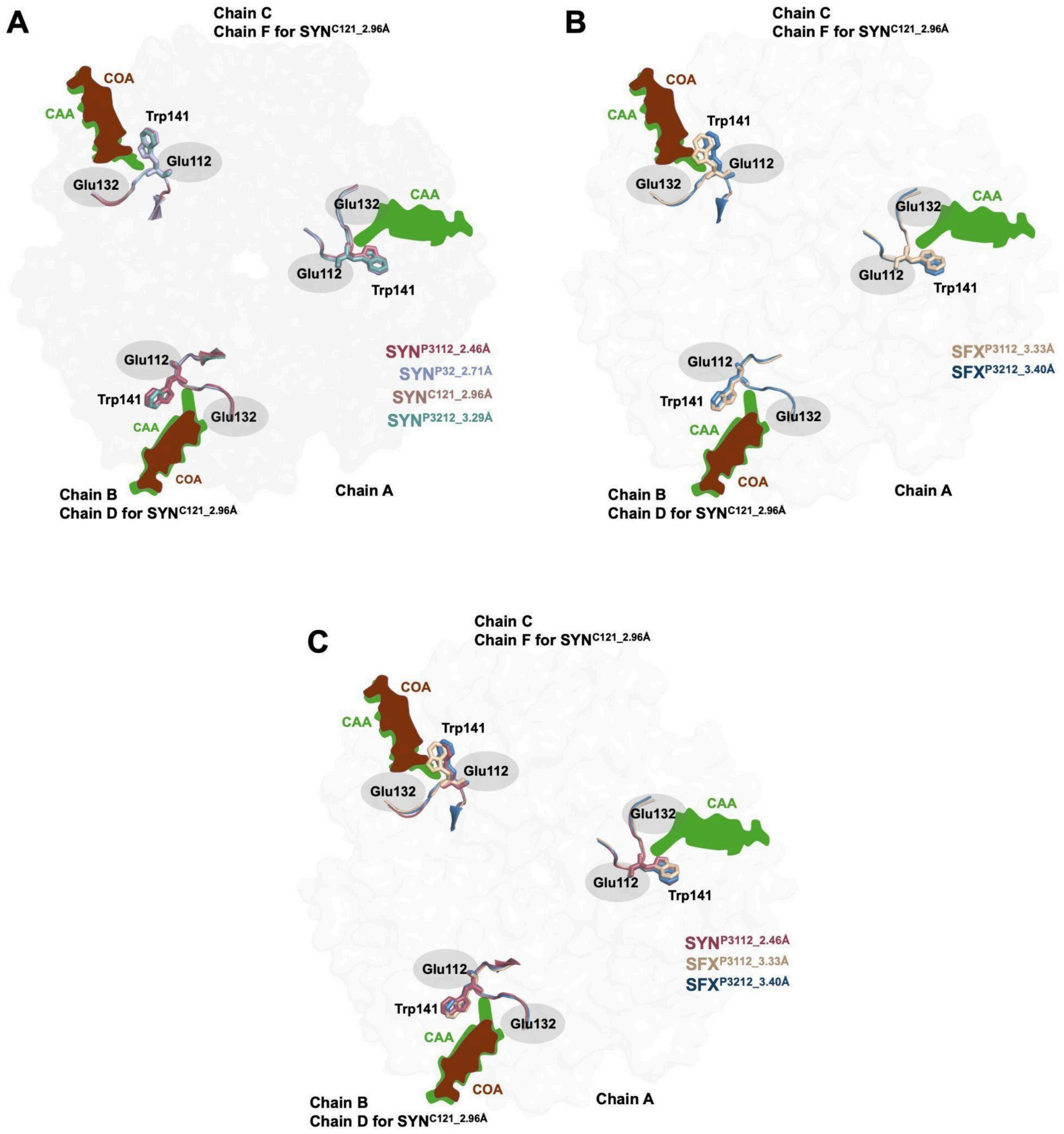


**Figure S10:** Representation of substrate entrance to the active site of the Nmar_1308 protein. **(A)** The critical position of the residue, Trp141, which limits the substrate size at the active site is shown for cryogenic temperature structures. **(B)** The critical position of the residue, Trp141, which limits the substrate (acetoacetyl-coenzyme A (CAA) and coenzyme A (COA)) size at the active site is shown for ambient temperature structures. **(C)** The residue Trp141 from SYN$^{P3112_2.46Å}$, SFX$^{P3112_3.33Å,}$ and SFX$^{P3212_3.40Å}$ structures are superposed to reveal the effect of minor side chain variations to enter the substrate.

**Table S1:** List of crystallization conditions.

| **SYN$^{P3112_2.46Å}$/ SFX$^{P3112_3.40Å}$** | **SYN$^{P32_2.71Å}$** | **SYN$^{C121_2.96Å}$** |
|---|---|---|
| 0.06 M Divalents (0.3M Magnesium chloride hexahydrate; 0.3M Calcium chloride dihydrate) | 0.09 M NPS (0.3M Sodium nitrate; 0.3 Sodium phosphate dibasic; 0.3M Ammonium sulfate) | 0.1 M Sodium chloride, 0.002 M Spermine tetrahydrochloride |
| 0.1 M Buffer System 3 (1.0 M Tris (base); BICINE pH 8.5) | 0.1 M Buffer System 3 (1.0 M Tris (base); BICINE pH 8.5) | Buffer system (0.05 M Bis-Tris pH 7.0) |
| 50% v/v Precipant Mix 1 (40% v/v PEG 500* MME; 20% w/v PEG 2000) | 37.5 % v/v Precipitant Mix 4 (25% v/v MPD; 25% PEG 1000; 25% w/v PEG 3350) | Precipitant (37 % w/v PEG 1000) |

| **SYN$^{P3212_3.29Å}$** | **SFX$^{P3112_3.33Å}$** |
|---|---|
| 0.09 M Hologens (0.3M Sodium fluoride; 0.3M Sodium bromide; 0.3 M Sodium iodide) | 0.06 M Divalents (0.3M Magnesium chloride hexahydrate; 0.3M Calcium chloride dihydrate)<br>0.1 M Buffer System 2 (1.0 M Tris (base); BICINE pH 8.5) |
| 0.1 M Buffer System 2 (1.0 M Tris (base); BICINE pH 8.5) | 0.1 M Buffer System 3 (1.0 M Tris (base); BICINE pH 8.5) |
| 37.5 % v/v Precipitant Mix 4 (25% v/v MPD; 25% PEG 1000; 25% w/v PEG 3350) | 50% v/v Precipitant Mix 4 (25% v/v MPD; 25% PEG 1000; 25% w/v PEG 3350) |
